# Two-dimensional NMR lineshape analysis of single, multiple, zero and double quantum correlation experiments

**DOI:** 10.1101/782631

**Authors:** Christopher A. Waudby, Margaux Ouvry, Ben Davis, John Christodoulou

**Affiliations:** Institute of Structural and Molecular Biology, University College London and Birkbeck College, London UK; Biotechnologie, Sanofi, Route d’Avignon, 30390 Aramon, France; Vernalis (R&D) Limited, Granta Park, Great Abington, UK

## Abstract

NMR spectroscopy provides a powerful approach for the characterisation of chemical exchange and molecular interactions by analysis of series of experiments acquired over the course of a titration measurement. The appearance of NMR resonances undergoing chemical exchange depends on the frequency difference relative to the rate of exchange, and in the case of one-dimensional experiments chemical exchange regimes are well established and well known. However, two-dimensional experiments present additional complexity, as at least one additional frequency difference must be considered. Here we provide a systematic classification of chemical exchange regimes in two-dimensional NMR spectra. We highlight important differences between exchange in HSQC and HMQC experiments, that on a practical level result in more severe exchange broadening in HMQC spectra, but show that complementary alternatives to the HMQC are available in the form of HZQC and HDQC experiments. We present the longitudinal relaxation optimised SOFAST-H(Z/D)QC experiment for the simultaneous acquisition of sensitivity-enhanced HZQC and HDQC spectra, and the longitudinal and transverse relaxation optimised BEST-ZQ-TROSY for analysis of large molecular weight systems. We describe the application of these experiments to the characterisation of the interaction between the Hsp90 N-terminal domain and a small molecule ligand, and show that the independent analysis of HSQC, HMQC, HZQC and HDQC experiments provides improved confidence in the fitted dissociation constant and dissociation rate. Joint analysis of such data may provide improved sensitivity to detect and analyse more complex multi-state interaction mechanisms such as induced fit or conformational selection.

## Introduction

The NMR lineshape (i.e. intensity, frequency, phase and linewidth) of a spin undergoing chemical exchange (i.e. a spin that is interconverting between multiple states in dynamic equilibrium) is well known to provide a powerful spectroscopic probe of the underlying exchange process. This lineshape may be modulated by a variety of means, including rf pulse sequences such as CPMG pulse trains^1^, *R*_1*ρ*_ spin locks^2, 3^ or saturation with a single frequency or frequency comb^4–6^; field shuttling^7^; or variation of external parameters such as protein or ligand concentration or temperature^8, 9^. The resulting modulations may then be fitted to determine details of the exchange process such as chemical shift differences, populations of states, and the rate of exchange between them. As such, NMR spectroscopy provide an indispensable, label-free tool for studying both intramolecular dynamics and biomolecular and other host-guest interactions^10–13^.

The effects of chemical exchange on lineshapes in one-dimensional NMR spectra are well understood^14^. Depending on the frequency difference, Δ*ω*, between two resonances, A and B, that are in exchange with rate *k*_ex_ = *k*_AB_ + *k*_BA_ (where *k*_AB_ and *k*_BA_ are the forward and backward rates respectively), slow and fast exchange regimes may be defined that have characteristic limiting behaviour. We do not distinguish in this work between the slow and slow-intermediate or fast and fast-intermediate regimes, which instead are encompassed within our definitions of the slow and fast exchange limits. We also assume that there is no significant difference in relaxation rates between states. In the slow exchange regime (*k*_ex_ ≪ |Δ*ω*|), resonance intensities are modulated according to their population (plus a lifetime line broadening contribution to the linewidth) but no chemical shift perturbations are observed (to first order in *k*_ex_). In contrast, in the fast exchange regime (*k*_ex_ ≫ |Δ*ω*|) a single resonance is observed at a population weighted average of the original chemical shifts, and with an exchange-induced contribution to the linewidth determined by both the frequency difference between states and the exchange rate (discussed further below). Lastly, when the exchange rate and frequency difference are comparable, severe line broadening can result (the ‘intermediate exchange’ regime). For equal populations of the exchanging states A and B, this is best characterised by the ‘coalescence point’ at which the two original resonances can no longer be distinguished (defined by the vanishing first and second derivatives of the lineshape), which occurs when 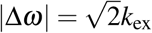^14^. Alternatively, exchange regimes may be characterised by the dependence of the exchange-induced line broadening term *R*_ex_ on the static magnetic field strength, *B*_0_, through the parameter *α* = *d*(ln *R*_ex_)*/d*(ln *B*_0_) which varies from 0 to 2 between the slow and fast exchange limits^15^.

As the appearance of one-dimensional NMR spectra can be modulated strongly by the chemical exchange process, observed spectra may be fitted quantitatively to numerical solutions of the Bloch-McConnell equations governing the evolution of magnetisation vectors (or at a more sophisticated level of theory, to the Liouville-von Neumann equation for the time evolution of the quantum mechanical density operator) in order to determine the microscopic rate constants for the exchange process, and the populations and chemical shifts of the various states^16, 17^. This procedure, termed ‘lineshape analysis’ or ‘dynamic NMR’, is particularly effective when a series of one-dimensional spectra can be analysed as a function of an external parameter such as temperature or ligand concentration, in order to determine parameters such as activation energies and dissociation constants.

The analysis of interactions within more complex macromolecules, such as proteins and nucleic acids, requires the use of two-dimensional NMR experiments such as the HSQC (heteronuclear single quantum correlation)^18^ or HMQC (heteronuclear multiple quantum correlation)^19^ to resolve the hundreds of resonances from component residues. While in some cases it is possible to perform one-dimensional lineshape analysis on cross-sections from such spectra^20, 21^, this approach risks a number of systematic errors^8^. Indeed, exchange of transverse magnetisation during preparation, chemical shift evolution and mixing periods has recently been shown to give rise to coherent cross-peaks that cannot be account for by one-dimensional treatments^9^. Instead, we have previously developed a truly two-dimensional lineshape analysis strategy, based on fitting observed two-dimensional spectra to complete quantum mechanical simulations of the underlying pulse sequence^8^. The analysis has been implemented in a software tool, TITAN, which has since been applied to study a wide range of biomolecular interactions^22–24^.

Two-dimensional lineshape analysis has two particular differences from the well-known one-dimensional theory. Firstly, there are now *two* chemical shift differences, Δ*δ*_*I*_ and Δ*δ*_*S*_, to consider and therefore it is not always possible to classify resonances as being simply in fast or slow exchange regimes. Secondly, in experiments such as the HMQC, chemical shift evolution occurs as a combination of zero and double quantum coherences, with frequencies *ω*_*S*_ *± ω*_*I*_, and it is the differences in these frequencies between states that should be compared to the exchange rate rather than the difference in single quantum frequencies. Indeed, we remarked in our original work on two-dimensional lineshape analysis that HSQC and HMQC experiments can give rise to different patterns of exchange broadening that must therefore be analysed with the appropriate framework^8^.

In this work, we expand on these earlier observations, by presenting a systematic classification of chemical exchange regimes appropriate for two-dimensional NMR experiments. We also consider zero- and double-quantum correlation experiments (HZQC and HDQC respectively), and describe new sensitivity-optimised pulse sequences for their acquisition that may offer a number of advantages over more commonly used HSQC and HMQC experiments.

## Results

### Classification of exchange regimes in two-dimensional NMR experiments

In this work, we consider heteronuclear correlation spectra involving two spins, labelled *I* and *S*, undergoing chemical exchange between two states, A and B, with rate *k*_ex_ = *k*_AB_ + *k*_BA_. We will also assume that the difference in relaxation rates between states is negligible relative to the frequency difference, i.e. Δ*R*_2_ ≪ |Δ*ω*|. It is then convenient to define the relative frequency differences *ξ*_*X*_ = Δ*ω*_*X*_ */k*_ex_, such that |*ξ* | ≪ 1 represents the fast exchange limit and | *ξ* | ≫ 1 represents the slow exchange limit (in a one-dimensional sense). The canonical coalesence point 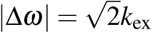 therefore corresponds to 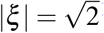^14^.

We will focus in particular on the analysis of molecular interactions using NMR titration measurements, exemplified by the interaction of a protein, P, with a ligand, L, to form a complex, PL:

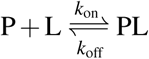

where the dissociation constant *K*_*d*_ = *k*_off_*/k*_on_. As the free ligand concentration, [L], always increases as ligand is added, the exchange rate, *k*_ex_ = *k*_on_[L] + *k*_off_, must also increase monotically with the fraction bound, *p*_*B*_, such that *k*_ex_ = *k*_off_*/*(1 − *p*_*B*_)^25^. The relative frequency difference, *ξ* = Δ*ω/k*_ex_ = (1 − *p*_*B*_)Δ*ω/k*_off_, therefore decreases as the titration proceeds, and so it is not always possible to classify a spin system as being in a single exchange regime throughout a titration. However, as the effects of chemical exchange on resonance lineshapes are most strongly manifest at the titration midpoint (*p*_*B*_ = 1*/*2), we will generally use the relative frequency differences at this point, *ξ* = Δ*ω/*2*k*_off_, as a convenient point of reference in discussions below.

### Single quantum correlation experiments

Single quantum (SQ) correlation experiments, such as the HSQC or (amide) TROSY experiments^18, 26^, have two characteristic parameters, *ξ*_*I*_ and *ξ*_*S*_, corresponding to the frequency differences in direct and indirect dimensions. Four two-dimensional exchange regimes may therefore be identified: fast exchange in both dimensions (FF, |*ξ*_*I*_| ≪ 1 and |*ξ*_*S*_| ≪ 1); slow exchange in both dimensions (SS, |*ξ*_*I*_| ≫ 1 and |*ξ*_*S*_| ≫ 1); fast exchange in the direct dimension and slow exchange in the indirect dimension (FS, |*ξ*_*I*_| ≪ 1 and |*ξ*_*S*_| ≫ 1); and lastly, slow exchange in the direct dimension and fast exchange in the indirect dimension (SF, |*ξ*_*I*_| ≫ 1 and |*ξ*_*S*_| ≪ 1). These regimes are marked schematically on a set of frequency axes in Fig. 1A. ‘Fast exchange’, in the sense of 2D resonances progressively changing frequency along the course of a titration, will only be observed within the ‘FF’ region where exchange is fast with respect to all relevant frequency differences.

**Figure 1.**
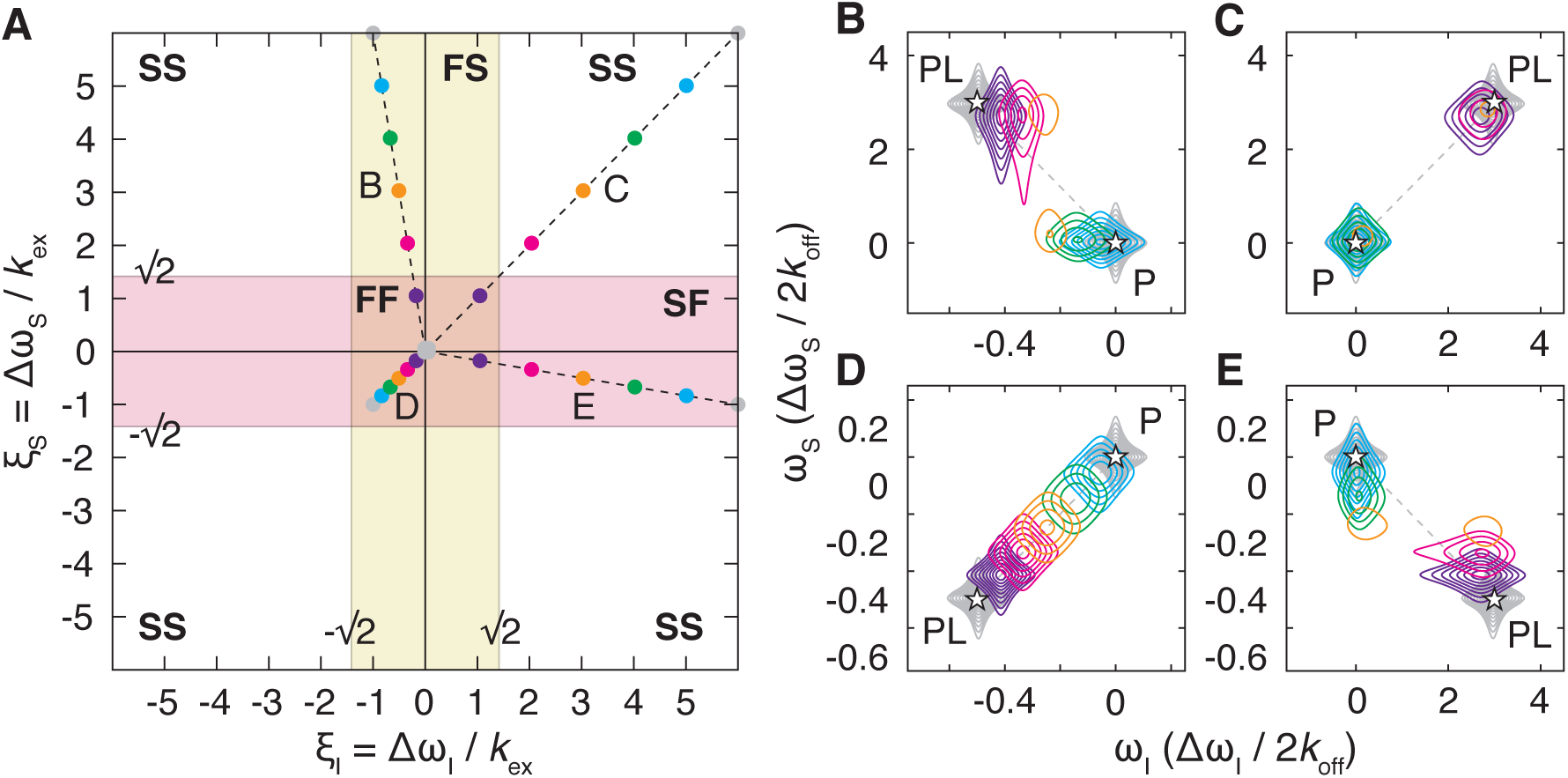
Classification and examples of two-dimensional exchange regimes for single-quantum correlation experiments. (A) Two-dimensional exchange regimes are plotted as a function of the relative frequency differences *ξ*_*I*_ = Δ*ω*_*I*_*/k*_ex_ and *ξ*_*S*_ = Δ*ω*_*S*_*/k*_ex_. A yellow band indicates the region of fast exchange in the direct dimension (bounded by the coalescence point), while the red band indicates the region of fast exchange in the indirect dimension. (B–E) Simulated titration measurements for residues having relative frequency differences (at the titration midpoint): (B) *ξ*_*I*_ = 0.5 and *ξ*_*S*_ = 3; (C) *ξ*_*I*_ = 3 and *ξ*_*S*_ = 3; (D) *ξ*_*I*_ = 0.5 and *ξ*_*S*_ = 0.5; and (E) *ξ*_*I*_ = 3 and *ξ*_*S*_ = 0.5. Asterisks mark the positions of the free and bound resonances. Ligand concentrations were selected so that the bound fraction varied uniformly from 0 to 99.9% (grey–blue–purple–grey). Contour levels are equal for all spectra, spaced at intervals incrementing 1.4x from a base value of 2% of the maximum intensity in the *apo* spectrum. The relative frequency differences, varying across the titration series, are plotted in panel A.

The observed perturbation Δ*ω*_obs_ to the frequency of the single resonance observed in the fast exchange limit, or to the frequency of the resonance of state A observed in the slow exchange limit, and the exchange contribution to the resonance linewidth, *R*_ex_, have been calculated for the indirect dimension to second order in *ξ* and *ξ*^−1^ for the fast and slow exchange limits respectively (see SI text). The resulting expressions recapitulate generally well known results^25^ and are presented in Table 1 for later comparison. In the fast exchange limit the observed frequency is simply the population weighted average of the individual states, while in the slow exchange limit a much smaller frequency perturbation is observed, that is only second order in the (relatively slow) exchange rate and inversely proportional to the (relatively large) frequency difference. Conversely, in fast exchange the exchange-induced line broadening is second order in the relatively small frequency difference, and inversely proportional to the relatively large exchange rate, and therefore will also generally be small in magnitude. The exchange contribution to the linewidth in slow exchange is determined by the lifetime line broadening, and is therefore more significant for the minor state where the populations are highly skewed. Corresponding relations for the direct dimension can be obtained by exchanging *S* and *I* terms. As the directly detected dimension is always a single quantum coherence, this is the same for all other experiments discussed below.

**Table 1.** Observed perturbations to the frequency, Δ*ω*, and linewidth, *R*_ex_, of the single resonance observed in the fast exchange limit, or of the resonance of state A in the slow exchange limit, calculated for single, multiple, zero and double quantum evolution periods.

| Exchange regime | Term | SQ | MQ | ZQ | DQ |
| --- | --- | --- | --- | --- | --- |
| Slow | $\Delta\omega_{obs}$ | $\frac{p_A p_B k_{ex}^2}{\Delta\omega_S}$ | $\frac{p_A p_B k_{ex}^2 \Delta\omega_S}{\Delta\omega_S^2 - \Delta\omega_I^2}$ | $\frac{p_A p_B k_{ex}^2}{\Delta\omega_S - \Delta\omega_I}$ | $\frac{p_A p_B k_{ex}^2}{\Delta\omega_S + \Delta\omega_I}$ |
| | $R_{ex}$ | $k_{AB}$ | $k_{AB}$ | $k_{AB}$ | $k_{AB}$ |
| Fast | $\Delta\omega_{obs}$ | $p_B \Delta\omega_S$ | $p_B \Delta\omega_S$ | $p_B (\Delta\omega_S - \Delta\omega_I)$ | $p_B (\Delta\omega_S + \Delta\omega_I)$ |
| | $R_{ex}$ | $\frac{p_A p_B \Delta\omega_S^2}{k_{ex}}$ | $\frac{p_A p_B (\Delta\omega_S^2 + \Delta\omega_I^2)}{k_{ex}}$ | $\frac{p_A p_B (\Delta\omega_S - \Delta\omega_I)^2}{k_{ex}}$ | $\frac{p_A p_B (\Delta\omega_S + \Delta\omega_I)^2}{k_{ex}}$ |

To illustrate these exchange regimes, a series of 2D spectra have been simulated according to a titration of a protein with a ligand, interacting through a simple two-state mechanism as discussed above. Ligand concentrations were chosen (relative to an arbitrary *K*_d_) to give bound populations varying uniformly from 0 to 99.9%. Chemical shift differences were defined according to the relative frequency difference at the titration midpoint, *ξ* = Δ*ω/*2*k*_off_, to represent the four 2D exchange regimes identified above. The simulated spectra are plotted in Fig. 1B–E, and the corresponding relative frequency differences, decreasing along the course of the titration, are marked on Fig. 1A. From these simulations, we observe that exchange in the ‘SS’ and ‘FF’ regimes (Fig. 1C and D respectively) appears as might be expected from 1D analogues. However, in the mixed ‘FS’ and ‘SF’ regimes more complex behaviour is observed, in which progressive chemical shift changes are observed along the ‘fast’ dimension only (Fig. 1B and E).

### Zero and double quantum correlation experiments

Having examined exchange regimes within the HSQC experiment, and as a prelude to the analysis of the more complicated HMQC experiment, we have considered exchange within zero and double quantum correlation experiments. Although not commonly acquired, amide zero quantum coherences have favourable relaxation rates, intermediate between single or multiple quantum coherences and the TROSY coherence *H*^*β*^*N*^*±*27^, while for methyl spin systems sharper lines can be obtained in HZQC spectra than in the standard ‘methyl-TROSY’ HMQC experiment itself^28^. Zero and double quantum correlation experiments have also been suggested as a means of alleviating exchange broadening in residues undergoing conformational exchange^29^.

The analysis of HZQC and HDQC experiments is essentially identical to that of the HSQC, with the difference in single quantum frequencies, *ξ*_*S*_, replaced by the difference in zero or double quantum frequencies, *ξ*_*S*_ *± ξ*_*I*_, as appropriate. Therefore, four two-dimensional exchange regimes can be identified (Fig. 2A,B) as described above for the HSQC (Fig. 1A). The observed frequency perturbations and exchange contribution to relaxation rates in the indirect dimensions have again been calculated (see SI texst) and are tabulated for slow and fast exchange limits in Table 1.

**Figure 2.**
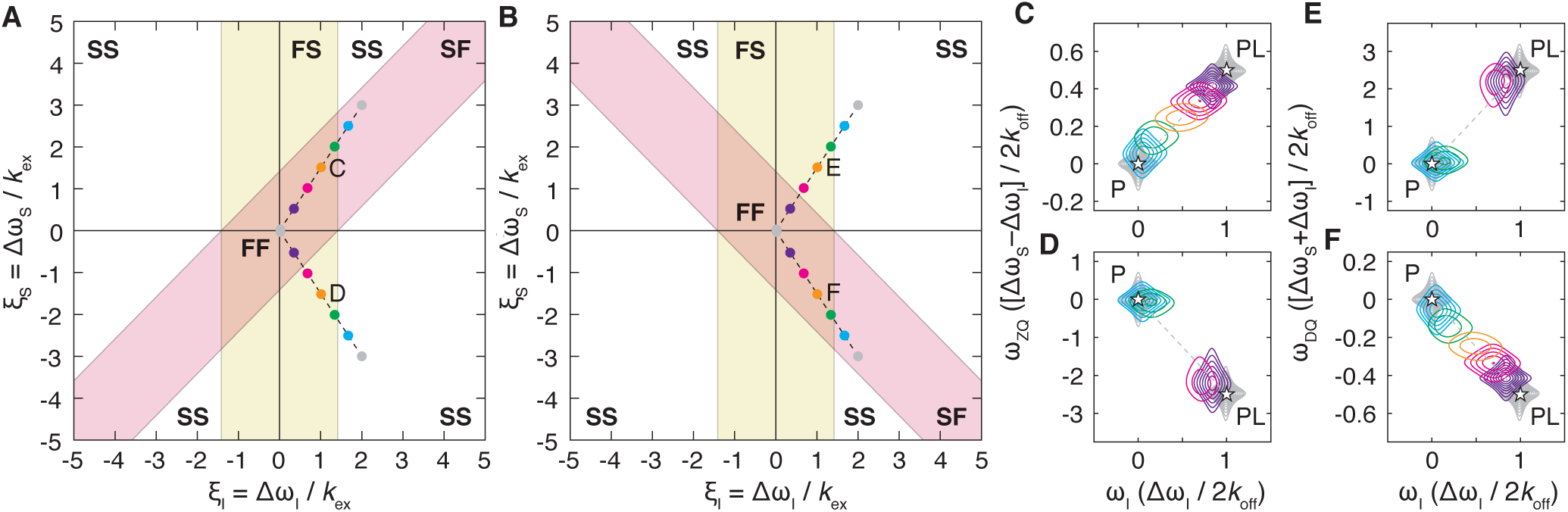
Classification and examples of two-dimensional exchange regimes for zero and double quantum correlation experiments. (A, B) Two-dimensional exchange regimes for (A) zero quantum and (B) double quantum correlation experiments are plotted as a function of the relative frequency differences *ξ*_*I*_ = Δ*ω*_*I*_*/k*_ex_ and *ξ*_*S*_ = Δ*ω*_*S*_*/k*_ex_. Yellow bands indicate the region of fast exchange in the direct dimension (bounded by the coalescence point), while red bands indicate the region of fast exchange in the indirect dimension, corresponding to the differences in zero or double quantum frequencies. (C–F) Simulated HZQC and HDQC titration measurements for residues having relative frequency differences (at the titration midpoint): (C,E) *ξ*_*I*_ = 1 and *ξ*_*S*_ = − 1.5; (D,F) *ξ*_*I*_ = 1 and *ξ*_*S*_ = − 1.5. Asterisks mark the positions of the free and bound resonances. Ligand concentrations were selected so that the bound fraction varied uniformly from 0 to 99.9% (grey–blue–purple–grey). Contour levels are equal for all spectra, spaced at intervals incrementing 1.4x from a base value of 2% of the maximum intensity in the *apo* spectrum. The relative frequency differences, varying across the titration series, are plotted in panels A and B.

It may be observed that the ranges of parameter space within the fast exchange (FF) regime for HZQC and HDQC experiments are complementary (Fig. 2A,B). This has two practical implications. Firstly, a resonance that is in the FS exchange regime within one experiment may be within the FF regime in the other, and this may facilitate tracking chemical shift changes during the course of a titration. This is illustrated for the simulated titration spectra shown in Fig. 2C–F. Secondly, due to the differences in exchange regimes between these experiments, two-dimensional lineshape analysis conducted on both sets of experiments should prove complementary, and their independent analysis may provide an additional layer of validation on results obtained from such fitting.

### Multiple quantum correlation experiments

In the HMQC experiment, magnetisation evolves as a mixture of zero and double quantum coherences during *t*_1_^19^. Thus, in contrast to the HSQC, there are three characteristic parameters to be considered (relative the to exchange rate): the frequency difference, *ξ*_*I*_, in the direct dimension, and the differences in both the zero and double quantum frequencies, *ξ*_*S*_ *± ξ*_*I*_, in the indirect dimension. From this, seven two-dimensional exchange regimes may therefore be identified, corresponding to fast and slow exchange in the direct dimension, and fast and slow exchange with respect to both zero and double quantum frequency differences, as indicated in Fig. 3A. ‘Fast exchange’ behaviour of 2D resonances will only be observed within the ‘FFF’ regime, in which exchange is fast with respect to all frequency differences. This is a smaller region of parameter space than for the single quantum experiment (Fig. 1A), with the practical consequence that during titrations it is likely to be more difficult to follow progressive chemical shift changes using HMQC experiments than using HSQC experiments.

**Figure 3.**
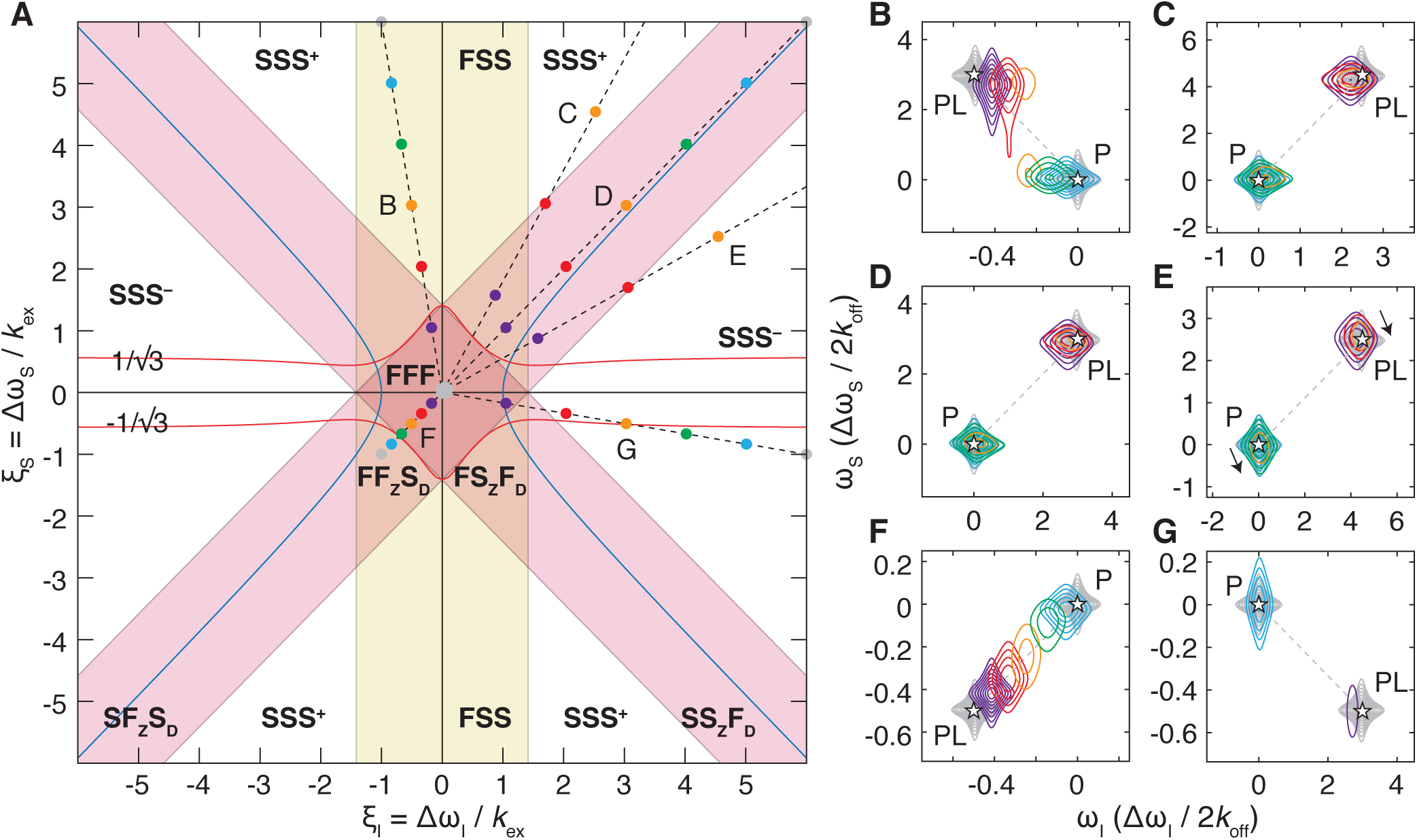
Classification and examples of two-dimensional exchange regimes for multiple-quantum correlation experiments. (A) Two-dimensional exchange regimes are plotted as a function of the relative frequency differences *ξ*_*I*_ = Δ*ω*_*I*_*/k*_ex_ and *ξ*_*S*_ = Δ*ω*_*S*_*/k*_ex_. A yellow band indicates the region of fast exchange in the direct dimension (bounded by the coalescence point), while red bands indicate the region of fast exchange in the indirect dimension, corresponding to zero and double quantum frequency differences. A red line indicates the multiple-quantum coalescence point (Eq. 1), and a blue line indicates the boundary at which the initial chemical shift change (i.e. when *p*_*B*_ ≪ 1) in the indirect dimension has the opposite sign to the true chemical shift difference. (B–G) Simulated titration measurements for residues having relative frequency differences (at the titration midpoint): (B) *ξ*_*I*_ = 0.5 and *ξ*_*S*_ = 3; (C) *ξ*_*I*_ = 3 and *ξ*_*S*_ = 3; (D) *ξ*_*I*_ = 4 and *ξ*_*S*_ = 3; (E) *ξ*_*I*_ = 0.5 and *ξ*_*S*_ = 0.5; (F) *ξ*_*I*_ = 3 and *ξ*_*S*_ = 0.5; and (G) *ξ*_*I*_ = 3 and *ξ*_*S*_ = 4. Asterisks mark the positions of the free and bound resonances. Ligand concentrations were selected so that the bound fraction varied uniformly from 0 to 99.9% (grey–blue–purple–grey). Contour levels are equal for all spectra, spaced at intervals incrementing 1.4x from a base value of 2% of the maximum intensity in the *apo* spectrum. The relative frequency differences, varying across the titration series, are plotted in panel A.

As for the HSQC experiment, the observed chemical shift perturbations and the exchange contribution to the resonance line broadening in the indirect dimension have been calculated for the slow and fast zero/double quantum exchange regimes and are tabulated in Table 1. The most significant difference between the HSQC and HMQC experiments is that chemical shift changes and line broadening in the *indirect* dimension now depend upon the frequency difference in the *direct* dimension also. Two observations are particularly striking. Firstly, the exchange contribution to line broadening, *R*_ex_, in the fast (F_Z_F_D_) exchange regime scales as 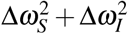, rather than 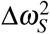 as in the single quantum case. Thus, HMQC resonances are subject to more significant line broadening than in the equivalent HSQC experiment. This may have practical consequences for the ability to detect and track chemical shift changes over the course of a titration experiment. Secondly, chemical shift perturbations in the slow (S_Z_S_D_) exchange regime are proportional to 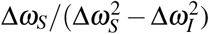. When the chemical shift difference in the direct dimension is large this term can become close to zero or even change sign, such that the initial direction of chemical shift changes over the course of a titration is hard to discern, or even occurs in the opposite direction to the bound peak position; we have labelled this regime ‘SS^−^’ (Fig. 3A). A calculation of chemical shift perturbations across all exchange regimes, in the limit *p*_*B*_ ≪ 1, indicates that this reversal occurs when 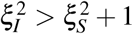. This region of parameter space is delineated with a blue line in Fig. 3B. This effect has previously been noted in the context of relaxation dispersion experiments, where the difference in chemical shifts of a resonance between HSQC and HMQC experiments can be used to infer the sign of the chemical shift difference to the unobserved excited state^30^.

It is also possible to calculate the coalescence point for the indirect dimension of the HMQC experiment. In contrast to the HSQC, for which the coalescence point occurs at 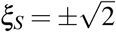, in the HMQC the coalescence point must be defined as a function of the frequency difference *ξ*_*I*_ also. By calculating the time evolution of the magnetisation analytically, including the central *πÎ*_*x*_ pulse to interconvert zero and double quantum coherences, and then taking the Fourier transform of the result, we obtain the lineshape in the indirect dimension, *y*(*ξ*_*S*_;*ξ*_*I*_). The coalescence point is then defined by a vanishing second derivative, 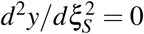, which yields:

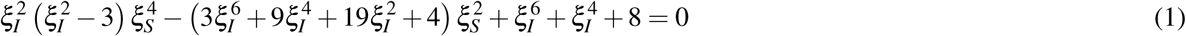

Solutions to this expression are plotted as red lines in Fig. 3A. Two further roots of Eq. 1 exist when |*ξ*_*S*_| ≳ 8, but inspection of the relevant lineshapes indicates that this does not correspond to coalescence behaviour but rather to distortions in the baseline between two well-resolved resonances. As expected, when *ξ*_*I*_ is zero then the coalescence point is identical to that of the HSQC, while for |*ξ*_*I*_| *<* 1 we observe that the region of parameter space below the coalescence point is well approximated by the requirement of being in fast exchange with respect to both zero and double quantum frequency differences (Fig. 3A). However, when |*ξ*_*I*_| becomes large then the coalescence point tends towards 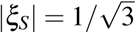. Moreover, direct inspection of simulated lineshapes within this regime 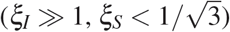 finds that, consistent with the limits calculated in Table 1, progressive chemical shift perturbations are not observed, and instead resonances interconvert with intensity modulations more similar to classical slow exchange behaviour (Fig. S1). Thus, for multiple quantum dimensions, we conclude that classical fast exchange behaviour is only observed within the F_Z_F_D_ regime.

To illustrate these cases further, a series of 2D titration spectra have been simulated (Fig. 3B–G), with relative frequency differences as indicated in Fig. 3A. In some cases, the simulated HMQC spectra are very similar to the equivalent HSQC spectra (Fig. 1). For example, HMQC spectra within the FSS regime (Fig. 3B), where *ξ*_*I*_ is small, are comparable to HSQC spectra within the FS regime (Fig. 1B). Exchange within the SSS^+^ and SF_Z_S_D_ regimes (Fig. 3C and D) is also similar, and comparable to that in the SS regime of HSQC spectra (Fig. 1C). However, in the SSS^−^ regime (Fig. 3E) we observe, as predicted above, that the direction of the small chemical shift perturbations in the indirect dimension is opposite to that of the exchanging state. A comparison of exchange in the FF regime of the HSQC (Fig. 1D) with the FFF regime of the HMQC (Fig. 1F) shows that the extent of line broadening is more severe in the HMQC. Close inspection also shows that non-linear chemical shift perturbations, which are often taken to be a sign of more complex association mechanisms, are observed in early stages of the titration (within the FF_Z_S_D_ regime). Lastly, resonances within the SF regime of the HSQC (Fig. 1E) are now within the SSS^−^ regime of the HMQC (Fig. 3G), close to the calculated coalescence point (Fig. 3A, red lines), and consequently very different spectra result, with severe line broadening but no chemical shift perturbations observed within the indirect dimension of the HMQC. In summary therefore, it is clear that more complex exchange phenomena are present within HMQC spectra, that on a practical level can result in stronger line broadening and altered chemical shift perturbations to those obtained through HSQC experiments.

### Longitudinal relaxation optimised pulse sequences for the measurement of heteronuclear zero and double quantum correlation spectra

Our analysis above of exchange within HZQC and HDQC spectra has indicated the potential utility of this pair of complementary experiments for the analysis of molecular interactions. A number of pulse sequences have been described for the measurement of ZQ and DQ correlation spectra. The first observation of ZQ and DQ transitions was reported by Müller in 1979^31^, and magnitude mode experiments were later developed in which separate ZQ and DQ transitions could be isolated^19^. It was also shown that ZQ and DQ transitions could be detected from the same physical experiment simply by applying different receiver phase cycling during processing, while experimental sensitivity could be enhanced using Ernst angle excitation^19^. The first phase sensitive HZQC and HDQC experiments were reported by Jarvet and Allard, including gradient selected variants^32^, and an alternative sensitivity-enhanced HZQC experiment was later reported by the Kay group for application to methyl groups^28^, with only a loss of 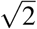 in sensitivity relative to the HMQC (and less when the improved relaxation of methyl ZQ coherences is taken into account).

Here we present further refinements of the sensitivity-enhanced HZQC experiment^28^. We have introduced longitudinal relaxation optimisation, applying the same principles as used for the SOFAST-HMQC experiment^33^. As established previously for the SOFAST-HMQC experiment, selective Ernst angle excitation of amide protons (or equally methyl protons if applied to a ^1^H,^13^C correlation experiment) is expected here also to provide a large sensitivity gain through efficient longitudinal cross-relaxation with the surrounding bath of unexcited spins and by avoiding saturation of the solvent, allowing rapid repetition rates. Alternatively, where sensitivity is not limiting, this enables rapid acquisition of the experiment.

The pulse sequence and coherence transfer pathways for the HZQC and HDQC experiments are shown in Fig. 4A. ^1^H +1 and −1 coherence orders are selected during *t*_1_ depending on the position of the 180° refocusing pulse indicated by dashed lines. Selection of Δ*p*_*S*_ = +1 or −1 is then required at the first 90° pulse on the S spin to isolate ZQ and DQ coherences. However, by storing individual steps of the phase cycle on this pulse and applying the required receiver phase during processing, both HZQC and HDQC experiments can be acquired simultaneously. Using the sensitivity enhanced approach, each of these spectra will have a factor of only 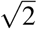 less sensitivity than original HMQC. Moreover, by shearing both ZQ and DQ spectra and then summing, a conventional correlation spectrum can in principle be obtained with identical sensitivity to the original HMQC (Fig. 4B).

**Figure 4.**
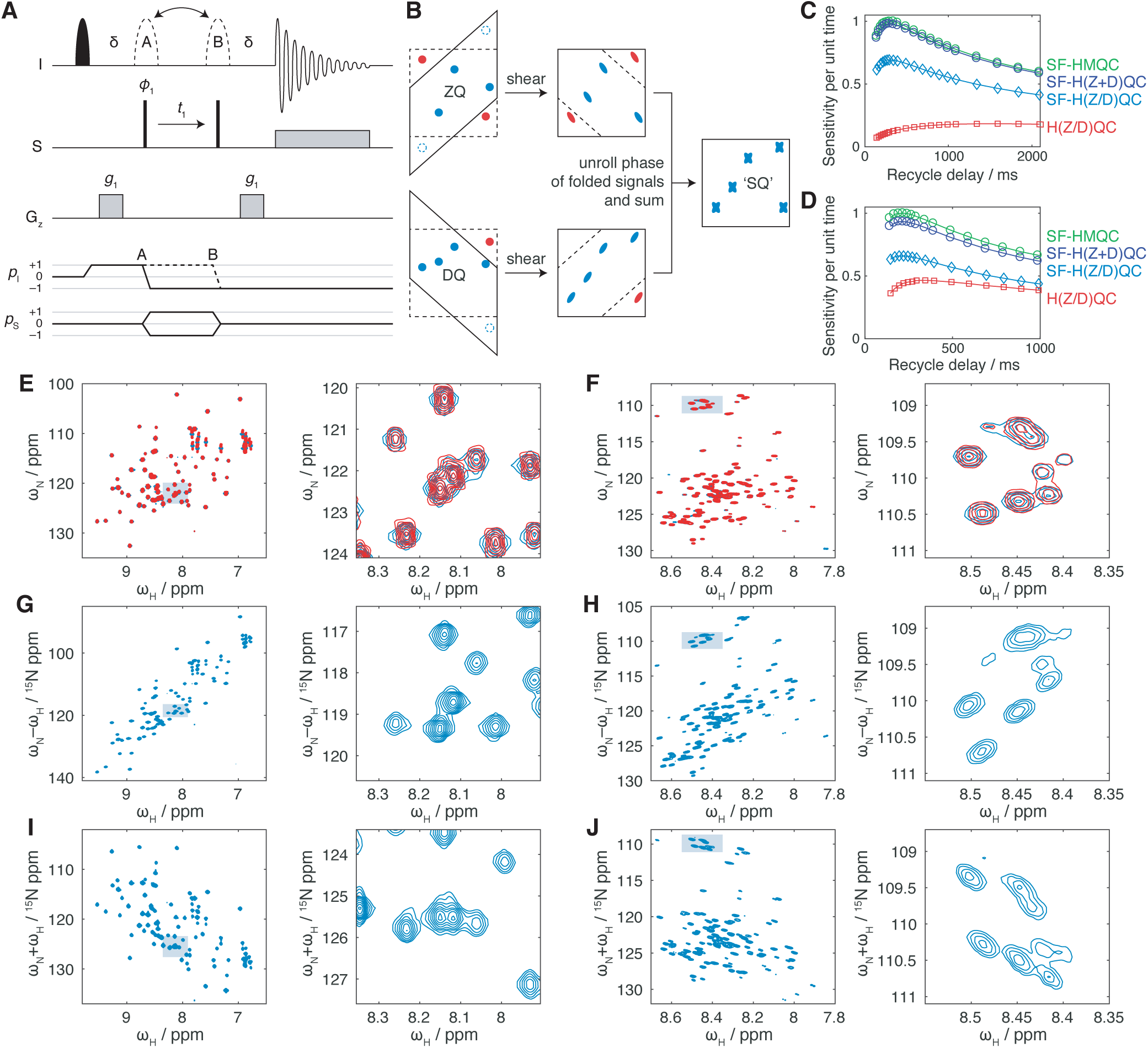
The SOFAST-H(Z/D)QC experiment. (A) Pulse sequence and coherence transfer pathways for the SOFAST-H(Z/D)QC experiment. Two experiments are recorded for each point in *t*_1_, in which the 180° pulse indicated with dashed lines is placed at positions A and B. The delay *δ* is 1*/*(2*J*) = 5.5 ms. The filled shaped pulse is a 120° Pc9 pulse (2657 *µ*s at 8.2 ppm, 700 MHz). The hollow pulses A and B are 180° Reburp pulses (1935 *µ*s at 8.2 ppm, 700 MHz). Phase cycling of *ϕ*_1_ is *x, − x* and the receiver phase *ϕ*_rx_ is *x, − x*. The phase of *ϕ*_1_ is incremented by 90° after each pair of experiments A and B, resulting in four FIDs labelled *A*_*x*_, *A*_*y*_, *B*_*x*_ and *B*_*y*_. Cosine and sine modulated components of ZQ and DQ spectra are formed from combinations of these FIDs: *ZQ*_cos_ = *A*_*x*_ + *iA*_*y*_ + *B*_*x*_ *−iB*_*y*_, *ZQ*_sin_ = *iA*_*x*_ *−A*_*y*_ *−iB*_*x*_ *−B*_*y*_, *DQ*_cos_ = *A*_*x*_ *−iA*_*y*_ + *B*_*x*_ + *iB*_*y*_, and *DQ*_sin_ = *− iA*_*x*_ *− A*_*y*_ + *iB*_*x*_ *− B*_*y*_. Note that spin dynamical phases are listed here, which when implemented may need to be adapted according to the spectrometer manufacturer^34, 35^. (B) Reconstruction of single-quantum-like H(Z+D)QC spectra by shearing and recombination of component ZQ and DQ spectra. (C, D) Relative sensitivity, per unit experimental time, measured from the integrated intensity of the first increment of the indicated experiments as a function of the recycle delay (including the acquisition period) for (C) ^1^H,^15^N-labelled ubiquitin (700 MHz, 298 K) and (D) ^1^H,^15^N-labelled FLN5 Y719E (700 MHz, 283 K). (E, F) SOFAST-HMQC (blue) and reconstructed SOFAST-H(Z+D)QC spectra (red) of (E) ^1^H,^15^N-labelled ubiquitin (700 MHz, 298 K) and (F) ^1^H,^15^N-labelled FLN5 Y719E (700 MHz, 283 K). Contour levels are matched between experiments, and the magnified region is indicated by the shaded box. (G–J) SOFAST-HZQC (G,H) and SOFAST-HDQC spectra (I,J) of (G,I) ^1^H,^15^N-labelled ubiquitin (700 MHz, 298 K) and (H,J) ^1^H,^15^N-labelled FLN5 Y719E (700 MHz, 283 K).

We have validated our new experiment, which we term the SOFAST-H(Z/D)QC, by comparison with ^1^H,^15^N SOFAST-HMQC measurements of the folded protein ubiquitin, and the unfolded Y719E variant of the FLN5 filamin domain^36^, 37. Analysis of the integrated amide signal intensity as a function of the inter-scan delay showed that for both samples the sensitivity of SOFAST-HZQC and SOFAST-HDQC experiments was very close to the expected factor of 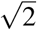 less than the SOFAST-HMQC (Fig. 4C,D). Very similar intensities were observed in the first increments of the SOFAST-HZQC and SOFAST-HDQC experiments and therefore only a single trace is shown in Fig. 4C,D. Comparison with an equivalent H(Z/D)QC experiment in which selective pulses in Fig. 4A were replaced with hard pulses shows that, as previously observed for the SOFAST-HMQC^33^, a substantial increase in sensitivity (or equivalently, a reduction in acquisition time) is obtained through the optimisation of longitudinal relaxation. This effect is particularly large in the case of the unfolded protein FLN5 Y719E (Fig. 4D), indicating the importance of avoiding saturation of solvent protons. Lastly, as expected, we find that the sensitivity of the combined ZQ and DQ experiments (Fig. 4B), which we term the SOFAST-H(Z+D)QC, is only marginally less than the original SOFAST-HMQC experiment (Fig. 4C,D).

SOFAST-HMQC, HZQC and HDQC spectra of ubiquitin are presented in Fig. 4E,G,I. Note that because ^15^N has a negative gyromagnetic ratio, the zero quantum frequency corresponds to the sum of the chemical shifts, *δ*_N_ + *δ*_H_, while the double quantum frequency corresponds to the difference in chemical shifts, *δ*_N_ −*δ*_H_. A larger spectral width was used to acquire these HZQC and HDQC experiments, but in practice this could be avoided as folded signals are unlikely to overlap with other resonances. As expected, HZQC and HDQC resonances are well resolved, with symmetric lineshapes. The sheared and reconstructed SOFAST-H(Z+D)QC spectrum (Fig. 4B) is also shown in red in Fig. 4E. This overlays closely with the SOFAST-HMQC spectrum, with comparable intensity. However, due to the shearing transform of the original Lorentzian tails in the direct dimensions of the ZQ and DQ spectra, the reconstructed cross-peaks have a discernible ‘X’ shape, with a slightly narrower linewidth on-resonance.

Similar spectra of the unfolded protein FLN5 Y719E are presented in Fig. 4F,H,J. In this case, due to the slower transverse relaxation associated with the more flexible polypeptide chain, the ^3^*J*_HNHA_ scalar coupling has a significant contribution to resonance linewidths in the direct dimension of the HMQC (Fig. 4F). Due to the amide selective refocusing pulse in the SOFAST-HMQC, this coupling is refocused during *t*_1_. However, in the SOFAST-HZQC and HDQC experiments the use of the selective pulse gives rise to an E.COSY effect^38^ that results in an undesirable diagonal appearance of resonances. This effect could be alleviated by deuteration of the protein at the H*α* position. Indeed, this has previously been reported to greatly increase the resolution and sensitivity of more conventional HSQC and HMQC measurements of intrinsically disordered proteins^39^.

### Experimental application to Hsp90 ligand binding

Having developed a theoretical understanding of the effects of chemical exchange on HSQC, HMQC, HZQC and HDQC experiments, and developed new sensitivity optimised pulse sequences for the measurement of HZQC and HDQC spectra, we sought to demonstrate their application to the analysis of a well-characterised protein–ligand interaction. For this, we selected the interaction of the 25 kDa N-terminal domain (NTD) of human Hsp90 (Fig. 5B) with the small molecule **1** (Fig. 5A), previously identified in a fragment screen^40^ and with a *K*_d_ of 42 *µ*M previously determined using surface plasmon resonance^41^. An NMR titration was carried out using 70 *µ*M uniformly ^1^H,^15^N-labelled Hsp90 NTD, and a series of fifteen HSQC, SOFAST-HMQC and SOFAST-H(Z/D)QC spectra were acquired at ligand concentrations from 0 to 800 *µ*M (Fig. 5C).

**Figure 5.**
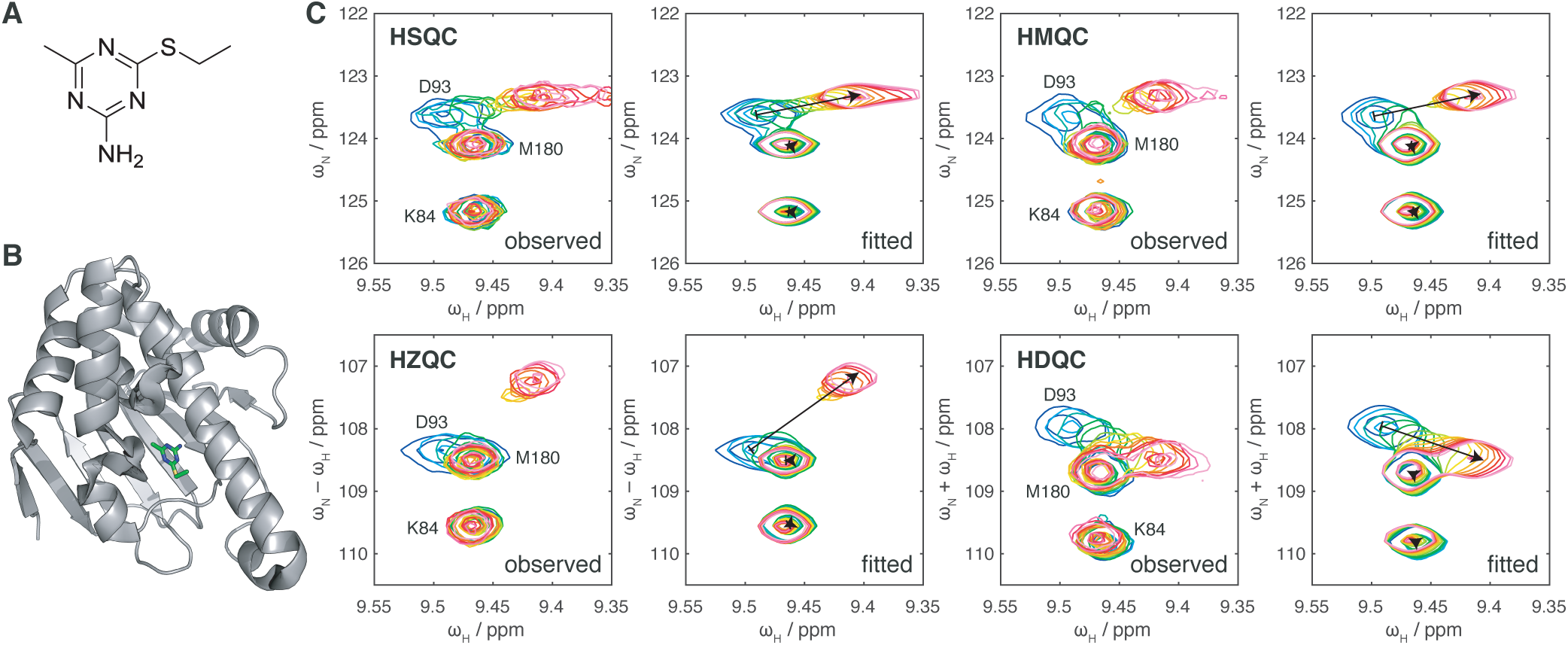
Measurement and two-dimensional lineshape analysis of the interaction between Hsp90 and a small molecule ligand. (A) Chemical structure of compound **1**^40, 41^. (B) Crystal structure of compound **1** in complex with human Hsp90 alpha N-terminal domain (pdb: 3b24)^41^. (C) Titration of 70 *µ*M ^1^H,^15^N human Hsp90 NTD with compound **1**, observed by HSQC, SOFAST-HMQC and SOFAST-H(Z/D)QC experiments as indicated, showing previously determined resonance assignments (298 K, 950 MHz)^42, 43^. Ligand concentrations were 0, 7.3, 14.5, 29, 51, 80, 116, 159, 209, 273, 350, 441, 552, 674 and 809 *µ*M (blue to red colouring). Two-dimensional fits are shown alongside with identical contour levels. Arrows indicate the fitted positions of free and bound resonances.

The resulting spectra illustrate experimentally the differences between the HSQC, HMQC, HZQC and HDQC experiments predicted above. This is particularly clear in the case of residue D93, which is depicted in Fig. 5C. This resonance has frequency differences Δ*ω*_*I*_ = 530 s^−1^ and Δ*ω*_*S*_ = 200 s^−1^ between the free and bound states, relative to a fitted dissociation rate (see below) of 787 *±* 19 s^−1^. All coherences for this residue are therefore formally in fast exchange regimes, but subject to varying amounts of exchange broadening (at the mid-point, *ξ*_*I*_ = 0.34, *ξ*_*S*_ = 0.13, *ξ*_*ZQ*_ = 0.46 and *ξ*_*DQ*_ = −0.21). Both HSQC and HDQC spectra show strong cross-peaks for this residue at all points in the titration, but in the HMQC and HZQC experiments exchange broadening is more severe.

Such variations in the appearance of a single residue across multiple experiments may be a useful tool to provide independent probes of chemical exchange, that will ultimately provide improved confidence in parameters such as dissociation rates that are obtained from analysis of titration data. To explore this, we performed independent two-dimensional lineshape analyses of each set of spectra, carrying out global fits of multiple residues exhibiting a range of exchange behaviours (Fig. 5C and S2–5). Uncertainties in the fitted parameters were evaluated using both the previously described block bootstrap resampling algorithm^8^, and jackknife resampling of spins or overlapping groups of spins (see Methods) (Table 2). Similar estimates of the dissociation constant and dissociation rate were obtained from all four sets of spectra. However, given the variation between individual fits, the bootstrap method appears to have underestimated the uncertainty in these parameters. The reason for this is not clear at present. Nevertheless, using the larger error estimates determined by the jackknife approach, consistent parameter estimates were obtained across all sets of measurements. The jackknife method has been implemented in v1.6 of TITAN and on the basis of these observations, we recommend that users perform error estimates using both approaches and choose the most conservative result. The combined dissociation constant determined from these measurements, 51.1 *±* 1.3 *µ*M, was slightly greater than the value of 42 *µ*M previously reported using SPR measurements^41^. This may reflect the effects of attachment to the surface or the biosensor, or the slightly lower ionic strength of the buffer used for SPR (50 mM Tris-based saline, pH 7.6, versus 50 mM sodium phosphate, 50 mM NaCl, pH 7.5).

**Table 2.** Hsp90 fit results. N.D., not determined. Uncertainties in fitted parameters calculated from two-dimensional lineshape analysis by block bootstrap resampling of residuals^8^ are indicated with a subscript *B*, while those obtained from jackknife sampling of multiple spins or spin groups (where overlapping) are indicated with a subscript *J*. The combined result shows the mean of all four NMR measurements, weighted according to the uncertainties determined by the jackknife method.

| Experiment | $K_d / \mu\text{M}$ | $k_{\text{off}} / \text{s}^{-1}$ |
| --- | --- | --- |
| SPR<sup>41</sup> | 42 | N.D. |
| HSQC | $45.5 \pm 0.7_B \pm 2.9_J$ | $835 \pm 24_B \pm 178_J$ |
| HMQC | $50.2 \pm 0.5_B \pm 2.5_J$ | $787 \pm 10_B \pm 31_J$ |
| HZQC | $53.3 \pm 1.1_B \pm 1.8_J$ | $779 \pm 24_B \pm 24_J$ |
| HDQC | $55.3 \pm 0.5_B \pm 6.0_J$ | $964 \pm 14_B \pm 122_J$ |
| Combined | $51.1 \pm 1.3$ | $787 \pm 19$ |

### Longitudinal and transverse relaxation optimised HZQC and HDQC pulse sequences

The SOFAST-H(Z/D)QC pulse sequence described above (Fig. 4A) is suitable for observations of amide resonances in intrinsically disordered proteins or folded domains with relatively low molecular weights (e.g. the 25 kDa Hsp90 NTD, Fig. 5). Perdeuteration will improve the accessible range of molecular weights further, but such systems are then most effectively observed using transverse relaxation optimised experiments, such as amide or methyl TROSY experiments. The amide SOFAST-H(Z/D)QC experiment (Fig. 4A) may be readily adapted and applied to methyl spin systems, in an analogous manner to the methyl-SOFAST-HMQC experiment^44^. The HZQC has already been reported to have more favourable relaxation properties than the ‘methyl-TROSY’ HMQC experiment^28^, and using the sequence described here an additional HDQC spectrum may be acquired at no cost. Additional variants have been reported incorporating filters for the fast-relaxing outer lines^45^, or in which heteronuclear polarisation may be used to enhance the sensitivity of HZQC experiments further^46^. However, two-dimensional lineshape analysis of spectra obtained using these more complex pulse sequences is not currently implemented in TITAN, and we therefore recommend the SOFAST-H(Z/D)QC sequence described above (Fig. 4A).

The case of amide spin systems is more complex. The TROSY experiment, in which the *H*^*β*^*N*^*±*^ transition is correlated with the *H*^−^*N*^*β*^ transition, has the most favourable transverse relaxation properties^27^. However, multiple quantum coherences also have a TROSY effect due to the absence of intra-system dipolar relaxation, and due to destructive CSA–CSA interference, ZQ coherences have particularly favourable relaxation properties. However, due to presence of transverse proton magnetisation, the relaxation of both ZQ and DQ coherences by external spins is more severe than for the *H*^*β*^*N*^*±*^ TROSY transition^26, 27^.

A sensitivity-enhanced ZQ-TROSY experiment has previously been described^27^. Here we present an updated BEST-ZQ-TROSY experiment that incorporates longitudinal relaxation optimisation using selective excitation of amide resonances (Fig. 6). In contrast to the SOFAST-H(Z/D)QC, it is not possible to acquire both ZQ and DQ experiments simultaneously, but a BEST-DQ-TROSY experiment may be acquired separately by modification of the pulse phases as described in the figure legend.

**Figure 6.**
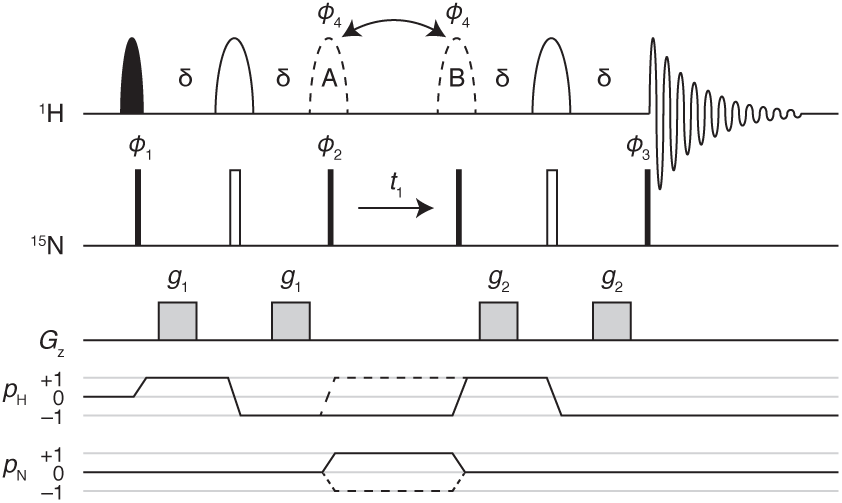
Pulse sequence and coherence transfer pathways for the BEST-ZQ-TROSY experiment. The filled shaped pulse is a 90° Pc9 pulse (1958 *µ*s at 8.2 ppm, 950 MHz) while the hollow pulses are 180° Reburp pulses (1432 *µ*s at 8.2 ppm, 950 MHz). The delay *δ* is 1*/*(4*J*) = 2.7 ms, *ϕ*_3_ is −*y, ϕ*_4_ is cycled *x*_4_, *y*_4_, (−*x*)_4_, (−*y*)_4_ and the receiver phase is cycled *x*, −*x*, −*y, y*, −*x, x, y*, −*y*. Gradients are applied as 1 ms trapezoidal pulses, *g*_1_ = 11% and *g*_2_ = 7%. Frequency discrimination is achieved by the echo/anti-echo method. The echo (dashed CTP) is acquired with the dashed pulse at ‘A’, *ϕ*_1_ −*x, x*, −*y, y* and *ϕ*_2_ −*y, y*, −*x, x*, and the anti-echo (solid CTP) is acquired with the dashed pulse at ‘B’, *ϕ*_1_ −*x, x, y*, −*y* and *ϕ*_2_ −*y, y, x*, −*x*. Alternatively, the BEST-DQ-TROSY experiment may be acquired by setting *ϕ*_3_ to *y*. The DQ echo is acquired with the dashed pulse at ‘A’, *ϕ*_1_ −*x, x*, −*y, y* and *ϕ*_2_ −*y, y, x*, −*x*, and the DQ anti-echo is acquired with the dashed pulse at ‘B’, *ϕ*_1_ −*x, x, y*, −*y* and *ϕ*_2_ −*y, y*, −*x, x*. Note that spin dynamical phases are listed here, which when implemented may need to be adapted according to the spectrometer manufacturer^34, 35^.

We have tested this new sequence experimentally with observations of ^2^H,^15^N-labelled ubiquitin in H_2_O at 277 K, 950 MHz (Fig. 7). While the rotational correlation time is still relatively low, ca. 9 ns, this nevertheless provides validation of the pulse sequence and points towards applications to higher molecular weight systems. Fig. 7A–D presents a comparison of BEST-TROSY^47^, BEST-HSQC^48^, BEST-ZQ-TROSY and BEST-DQ-TROSY experiments, which were all acquired with identical acquisition times and parameters. Folded resonances can be observed in both ZQ and DQ experiments (magenta contours), but it is clear that this does not introduce additional resonance overlap. Cross-sections through the representative E64 resonance are shown in Fig. 7E–H, and these were fitted to Lorentzian lineshapes to determine the indicated linewidths. As expected, TROSY, ZQ-TROSY and DQ-TROSY experiments all had reduced linewidths in the direct dimension relative to the HSQC. In the indirect dimension, the ZQ resonance was marginally sharper than the HSQC resonance, and approximately 1.6 times broader than the TROSY, while the DQ resonance was substantially more broad (2.6 times that of the TROSY). However, the intensities of the TROSY and ZQ-TROSY resonances were comparable, and significantly higher than in both HSQC and DQ-TROSY spectra. Thus, we believe the BEST-ZQ-TROSY experiment may be a useful and reasonably sensitive experiment for applications in high molecular weight systems. Importantly, because coherence transfer in this experiment is much simpler than in the regular TROSY experiment (in which the back-transfer *H*^*β*^*N*^*±*^ → *H*^−^*N*^*β*^ occurs through several stages), the ZQ-TROSY experiment (and indeed the DQ-TROSY experiment) is amenable to two-dimensional lineshape analysis.

**Figure 7.**
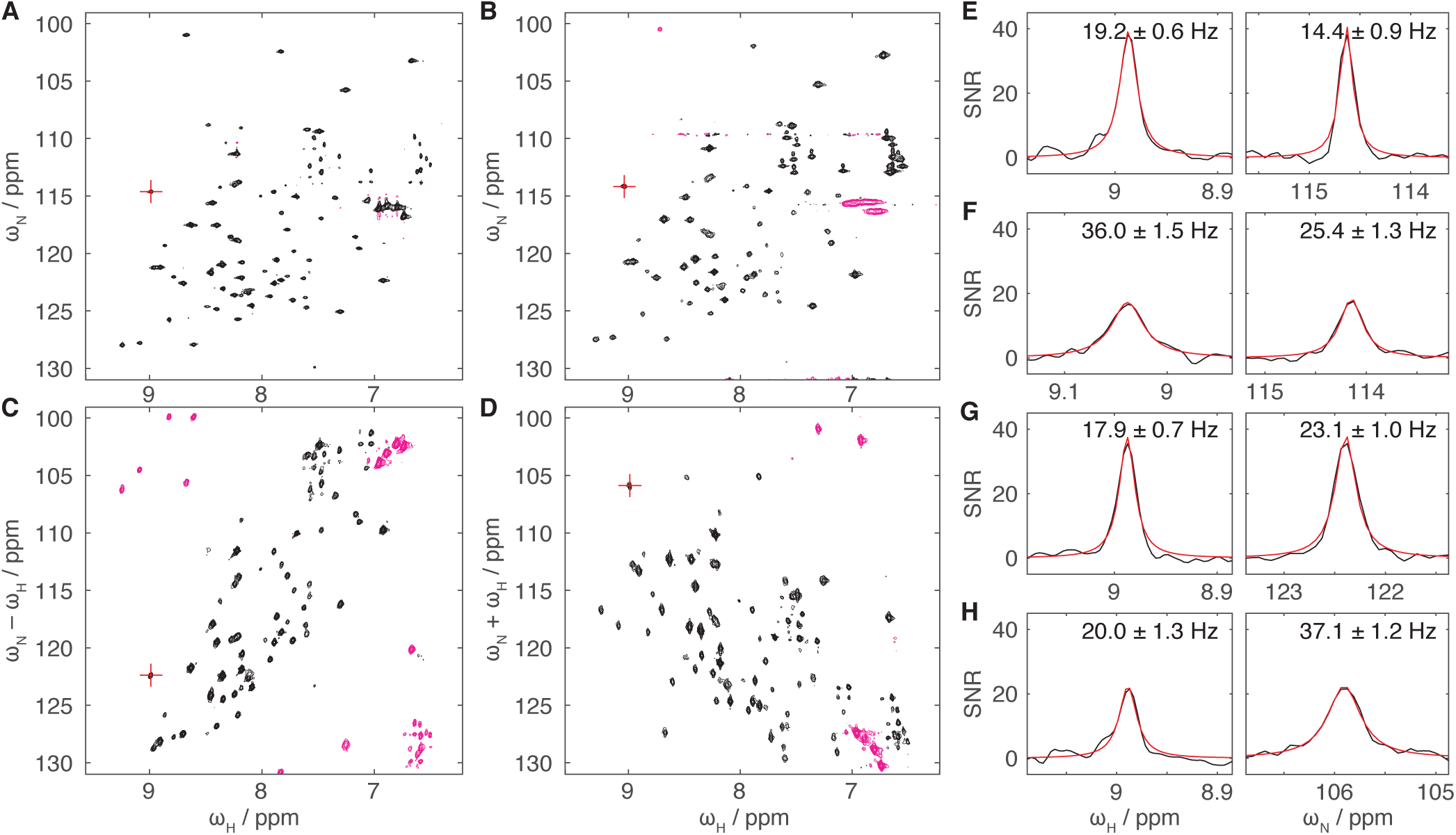
(A) BEST-TROSY, (B) BEST-HSQC, (C) BEST-ZQ-TROSY and (D) BEST-DQ-TROSY spectra of ^2^H,^15^N-labelled ubiquitin in H_2_O (277 K, 950 MHz). Spectra were acquired with identical acquisition times and are shown with matched contour levels (magenta contours indicate negative intensities). Cross-sections through the E64 resonance, indicated with red crosshairs in panels A–D, are plotted for (E) BEST-TROSY, (F) BEST-HSQC, (G) BEST-ZQ-TROSY and (H) BEST-DQ-TROSY spectra. Red lines show fits to Lorentzian lineshapes with linewidths as indicated.

## Discussion

In this paper we have presented a systematic analysis of chemical exchange regimes within two-dimensional correlation spectra, focusing in particular on differences between SQ, MQ, ZQ and DQ-based experiments. In many ways, these results are not new. We have previously commented upon the differences in exchange behaviour between HSQC and HMQC experiments^8^, and indeed the use of ZQ and DQ TROSY experiments to reduce chemical exchange-induced resonance broadening has also been suggested previously^29^. Differences in ZQ and DQ relaxation rates have been proposed and exploited as a simple way to identify and characterise chemical exchange^49–52^, and comparisons between chemical shift perturbations in HSQC and HMQC experiments have been used to determine the signs of chemical shift differences in relaxation dispersion measurements^30^. Indeed, joint analysis of (exchange-induced) relaxation dispersion of SQ, MQ, ZQ and DQ coherences has been a powerful tool for the analysis of sparsely populated intermediates^53–55^. However, we believe that this is the first time that these results have been set out comprehensively, and in particular that the impact on chemical exchange regimes across NMR titration measurements has been fully considered.

A key practical result of our analysis is that HMQC experiments are much more sensitive to chemical exchange-induced line broadening than their HSQC (and amide TROSY) counterparts. Such sensitivity may be useful in some circumstances, but in others it may be a hindrance to tracking the movement of resonances across a series of titration measurements. As an alternative or complement, we have therefore developed the SOFAST-H(Z/D)QC experiment (Fig. 4A) for the simultaneous and sensitive acquisition of ZQ and DQ correlation spectra. Such spectra have reduced sensitivity to chemical exchange (Fig. 2) compared to the original HMQC, and this may be particularly useful in the case of titrations observed using methyl labelling. We have also developed longitudinal relaxation optimised BEST-ZQ-TROSY and BEST-DQ-TROSY experiments for application to amide spin systems (Fig. 6).

While transverse relaxation of ZQ coherences is not quite as optimal as in the full TROSY experiment (Fig. 7), the simpler pulse sequence means that these experiments are amenable to two-dimensional lineshape analysis, in contrast to standard TROSY experiments.

We have explored the difference between SQ, MQ, ZQ and DQ correlation experiments on experimental titration measurements by observing the interaction of the Hsp90 NTD with a small molecule ligand (Fig. 5)^41^. As expected, resonances with the HSQC, HMQC, HZQC and HDQC experiments that were acquired exhibited different chemical exchange behaviours, and the independent two-dimensional lineshape analysis of the four titration series using TITAN ultimately provided improved confidence in the dissociation constant and dissociation rate determined (Table 2). As the additional time required to acquire a second correlation spectrum during a titration measurement is often small relative to the total time required for setup and sample handling, we suggest that this may be a useful strategy in general: either by acquiring HSQC and HMQC measurements sequentially, or acquiring HZQC and HDQC experiments simultaneously as described above using the SOFAST-H(Z/D)QC experiment.

An additional virtue of such an approach is that it provides increased sensitivity to departures from simple two-state association mechanisms. NMR lineshapes are well known to be sensitive probes of more complex binding mechanisms, such as induced fit or conformational selection^8, 56, 57^, and the independent analysis of multiple experiment series may reduce the chance of overlooking such mechanisms. Equally, where such multi-state mechanisms have been identified, the analysis of multiple experiments may provide increased confidence in the fitted parameters. Ultimately, in analogy to analyses of relaxation dispersion measurements^53–55^, we envisage the global analysis of multiple experiment types (or indeed experiments at multiple field strengths), and efforts are underway to implement this in a future version of TITAN.

## Methods

### Sample preparation

Hsp90 protein was produced as described previously^58^. The N-terminal fragment of Hsp90alpha (residues 9– 236) was overexpressed in the E. Coli strain BL21 (pLysS) grown in EnPresso defined media (SigmaAldrich) supplemented with 4 g/L ^15^N ammonium chloride. The construct included a deca-his tag within a 15 amino acid N-terminal extension of MGHHHHHHHHHHSSGH, and the protein was purified using a Ni^2+^ affinity column followed by a monoQ ion-exchange column.

Compound **1** was identified from a fragment screen against Hsp90 as described previously^40^.

### NMR spectroscopy

#### Acquisition and processing

All NMR data were acquired using Bruker spectrometers operating with Topspin Pulse sequences for the SOFAST-H(Z/D)QC and BEST-ZQ-TROSY experiments are provided in supporting information. NMR data were processed using nmrPipe^59^. SOFAST-H(Z+D)QC spectra were sheared and recombined in nmrPipe using the shear.M macro. Scripts and additional processing macros for the analysis of the SOFAST-H(Z/D)QC experiment are provided in supporting information.

#### Validation of the SOFAST-H(Z/D)QC experiment

NMR experiments were acquired at 700 MHz to validate and assess the sensitivity of the SOFAST-H(Z/D)QC experiment using a ca. 1 mM sample of ^1^H,^15^N-labelled ubiquitin (10% D_2_O, 298 K) and a 350 *µ*M sample of ^1^H,^15^N-labelled FLN5 Y719E (10% D_2_O, 283 K). SOFAST-HMQC experiments^33^ were acquired using the same shaped pulses as specified for the SOFAST-H(Z/D)QC (Fig. 4A) while control measurements were recorded replacing shaped pulses with corresponding hard pulses. Sensitivity was assessed from the integrated amide intensity in the first increment of experiments recorded with a constant number of scans and a 50 ms acquisition time, and recycle delays varied from 50 ms to 2 s. Measurements were normalised for the total acquisition time (excluding dummy scans). 2D correlation spectra were acquired with a 100 ms recycle delay, a 100 ms acquisition time (1024 complex points) in the direct dimension and a 100 ms acquisition time (512 complex points) in the indirect dimension.

#### Hsp90 titration measurements

A 70 *µ*M sample of ^1^H,^15^N-labelled human Hsp90 N-terminal domain (50 mM sodium phosphate, 50 mM NaCl, pH 7.5, 10% D_2_O) was titrated with a 20 mM stock of ligand **1** (Fig. 5A) in d_6_-DMSO. 15 points were recorded, up to a maximum ligand concentation of 810 *µ*M. NMR experiments were acquired at 298 K, 950 MHz, with a 100 ms acquisition time (1536 complex points) in the direct dimension and a 21 ms acquisition time (64 complex points) in indirect dimensions. SOFAST-HMQC^33^ and SOFAST-H(Z/D)QC experiments were acquired with a recycle delay of 300 ms, and FHSQC experiments^60^ were acquired with a 1 s recycle delay.

#### BEST-ZQ-TROSY and BEST-DQ-TROSY experiments

NMR experiments were acquired at 950 MHz to validate and assess the sensitivity of the BEST-ZQ-TROSY and BEST-DQ-TROSY experiments using a ca. 1 mM sample of ^2^H,^15^N-labelled ubiquitin (10% D_2_O, 277 K). BEST-TROSY spectra^47^ were acquired using the b_trosyf3gpph.2 library pulse sequence. Spectra were acquired with a 100 ms recycle delay, a 100 ms acquisition time (1536 complex points) in the direct dimension and an 83 ms acquisition time (256 complex points) in the indirect dimension.

#### Lineshape analysis

Two-dimensional lineshape fitting, and simulations of HSQC, HMQC, HZQC and HDQC spectra, were carried out using TITAN (v1.6)^8^. HZQC and HDQC experiments have been implemented by a simple modification of the original HMQC implementation^8^ to isolate the subspace of ZQ or DQ coherences. Fitting of the Hsp90 titration data was carried out in two stages. The chemical shifts and linewidths of the unbound state were initially fitted using only the first spectrum, acquired in the absence of ligand. These parameters were then fixed, and the chemical shifts and linewidths of the bound state were fitted together with the dissociation constant and the dissociation rate. Bootstrap error estimates were calculated using 100 replicas. A second estimate of uncertainty in the dissociation constant and dissociation rate was also determined using a jackknife algorithm, in which fitting was repeated multiple times, with each spin system being omitted in turn. The reported uncertainties from this method are the standard deviation across all fits. The jackknife algorithm has been implemented in the v1.6 release of TITAN.

## Acknowledgements

We thank Anais Cassaignau for providing the sample of FLN5 Y719E, and Frank Delaglio for help preparing nmrPipe macros. We acknowledge the use of the UCL Biomolecular NMR Centre and the staff for their support. This work was supported by the Francis Crick Institute through provision of access to the MRC Biomedical NMR Centre. The Francis Crick Institute receives its core funding from Cancer Research UK (FC001029), the UK Medical Research Council (FC001029), and the Wellcome Trust (FC001029). This work was supported by a Wellcome Trust Investigator Award (to J.C., 206409/Z/17/Z).

## Additional information

Fig. S1: Simulated one-dimensional lineshapes for exchange in single, multiple, zero and double quantum frequency dimensions. Fig. S2–5: Observed and fitted HSQC, HMQC, HZQC and HDQC spectra of Hsp90 titration with compound **1**. Listings S1–7: Pulse sequences, parameter sets, processing scripts and processing macros for the SOFAST-H(Z/D)QC and BEST-ZQ-TROSY experiments.

