## Supplementary Information for "Two-dimensional NMR lineshape analysis of single, multiple, zero and double quantum correlation experiments"

**This PDF file includes:**

Supplementary Text
**Fig. S1:** Simulated one-dimensional lineshapes for exchange in single, multiple, zero and double quantum frequency dimensions.
**Figs. S2 to S5:** Observed and fitted HSQC, HMQC, HZQC and HDQC spectra of Hsp90 titration with compound **1**
**Listing S1:** SOFAST-H(Z/D)QC pulse program
**Listing S2:** SOFAST-H(Z/D)QC parameter file
**Listing S3:** SOFAST-H(Z/D)QC nmrPipe processing template
**Listing S4:** shufZQ.m (nmrPipe processing macro)
**Listing S5:** shufDQ.m (nmrPipe processing macro)
**Listing S6:** BEST-ZQ/DQ-TROSY pulse program
**Listing S7:** Parameter file for BEST-ZQ-TROSY

**Supplementary Text**

Calculation of chemical shift perturbations and exchange broadening

Evolution of single, zero or double quantum magnetisation $\boldsymbol{M} = (M_A \quad M_B)'$, in the presence of chemical exchange between states *A* and *B* is described, in the rotating frame of spin *A*, by the Liouvillian, *L*:

$$L = \begin{pmatrix} -k_{ex}p_B - R & k_{ex}p_A \\ k_{ex}p_B & -k_{ex}p_A - R + i\,\Delta\omega \end{pmatrix}$$

where the transverse relaxation rates, *R*, are assumed to be equal, and the frequency difference $\Delta\omega = \omega_S^B - \omega_S^A$ in the case of single quantum magnetisation, $\Delta\omega = (\omega_S^B - \omega_I^B) - (\omega_S^A - \omega_I^A)$ in the case of zero quantum magnetisation, and $\Delta\omega = (\omega_S^B + \omega_I^B) - (\omega_S^A + \omega_I^A)$ in the case of double quantum magnetisation. Without loss of generality, the relaxation rate *R* may be factored out, as it will contribute a constant amount to all resonances. The frequency and exchange contribution to the linewidth of the observed resonances are then given by the imaginary and real parts of the eigenvalues of *L* respectively.

In the fast exchange limit ($\Delta\omega \ll k_{ex}$) we may write *L* in terms of the dimensionless frequency difference $\xi = \Delta\omega/k_{ex}$, eliminating $p_A$ and choosing units of time such that $k_{ex} = 1$:

$$L_F = \begin{pmatrix} -p_B & 1 - p_B \\ p_B & -1 + p_B + i\,\xi \end{pmatrix}$$

The eigenvalues of $L_F$ are $-\frac{1}{2}\left[1 - i\,\xi \pm \sqrt{(1 - i\,\xi)^2 + 4i\,p_B\xi}\right]$. A series expansion of the principle eigenvalue, $-\frac{1}{2}\left[1 - i\,\xi - \sqrt{(1 - i\,\xi)^2 + 4i\,p_B\xi}\right]$, to second order in $\xi$ then provides the required results upon restoring $\Delta\omega$ and $k_{ex}$ (Table 1).

Similarly, in the slow exchange limit ($k_{ex} \ll \Delta\omega$) we may write *L* in terms of the dimensionless exchange rate $\rho = \mathrm{k_{ex}}/\Delta\omega$, eliminating $p_A$ and choosing units of time such that $\Delta\omega = 1$:

$$L_S = \begin{pmatrix} -p_B\rho & (1 - p_B)\rho \\ p_B\rho & -(1 - p_B)\rho + i \end{pmatrix}$$

The eigenvalues of $L_S$ are $-\frac{1}{2}[\rho - i \pm \sqrt{-1 + \rho(\rho + 4ip_B - 2i)}]$, and a series expansion to second order in $\rho$ then provides the required results upon restoring $\Delta\omega$ and $k_{ex}$ (Table 1).

In the case of multiple quantum evolution, magnetisation is interconverted between zero and double quantum coherences at the midpoint of the evolution period. The observed frequency perturbation and exchange broadening is then an average of that calculated above for zero and double quantum coherences, and the results of this averaging are presented in Table 1 for fast exchange with respect to both ZQ and DQ frequency differences, and slow exchange with respect to both ZQ and DQ frequency differences.

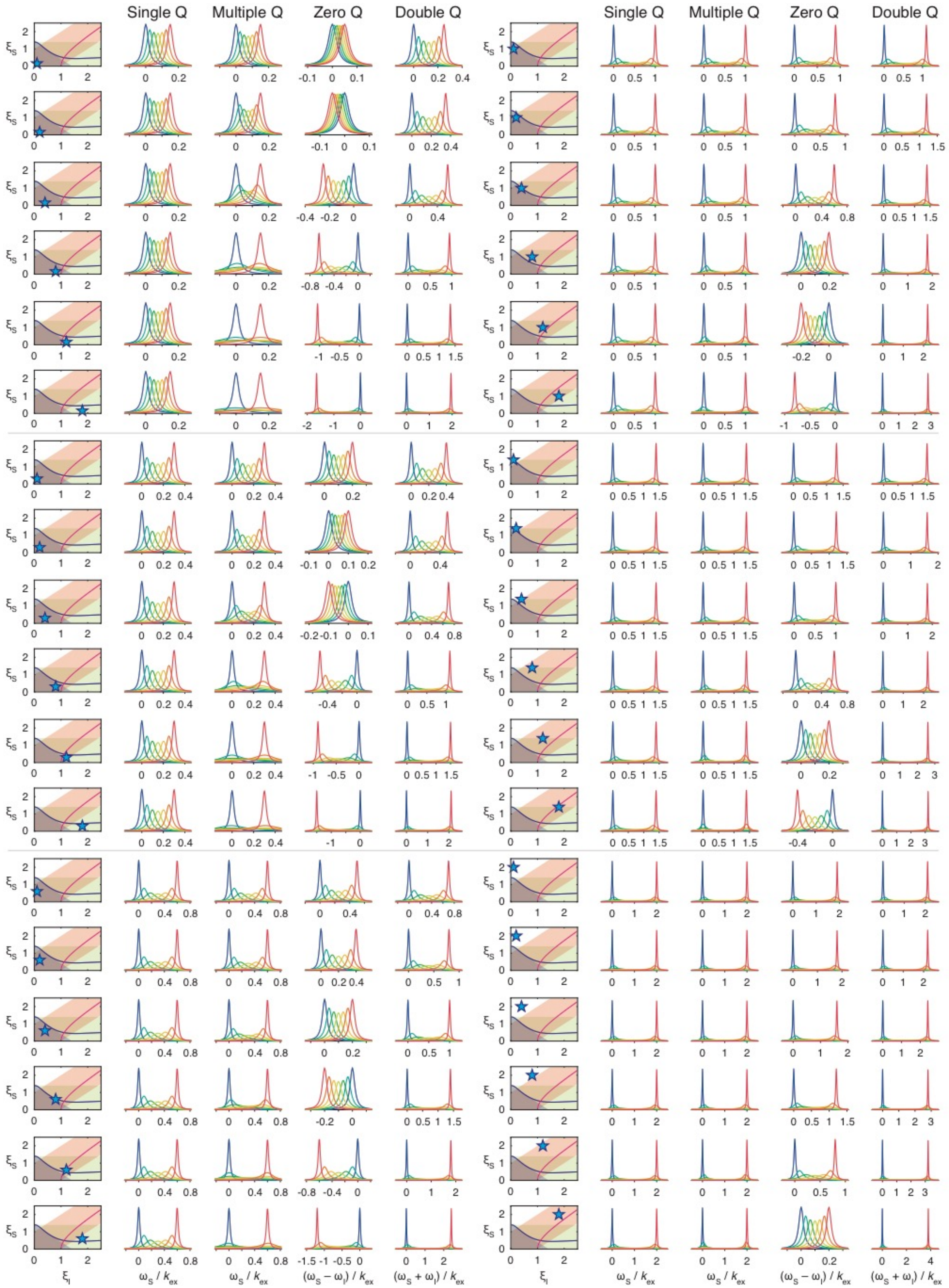


**Fig. S1.** Simulated one-dimensional lineshapes for $A \rightleftharpoons B$ exchange in single, multiple, zero and double quantum frequency dimensions. Lineshapes are calculated for populations of state B linearly spaced from 0 to 100% (blue to red colouring), normalised chemical shift differences, $\xi_{\mathrm{I}} = \Delta\omega_{\mathrm{I}}/k_{\mathrm{ex}}$ and $\xi_{\mathrm{S}} = \Delta\omega_{\mathrm{S}}/k_{\mathrm{ex}}$ as indicated (blue star in LH panels), and transverse relaxation rates $R_2 = k_{ex}/50$. 'Fast exchange' regimes (i.e. below the coalescence point) are indicated with green shading (single quantum), red shading (zero quantum) and blue shading (double quantum), while the multiple quantum coalescence point is indicated with a solid blue line (LH panels). A solid magenta line (LH panels) indicates the point at which the initial chemical shift change (i.e. when $p_B \ll 1$) for multiple quantum dimensions changes sign.

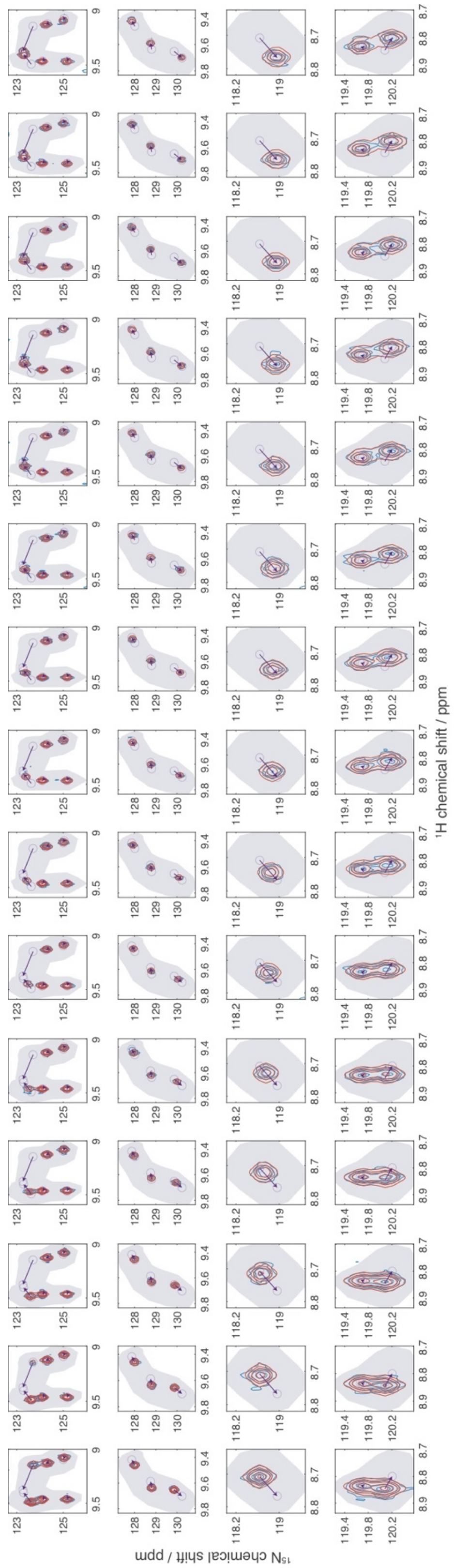


**Fig. S2.** $^{1}$H,$^{15}$N HSQC measurements (blue) and TITAN fits (red) of Hsp90 titration with compound **1** (Fig. 5). Grey shading indicates fitted ROIs.

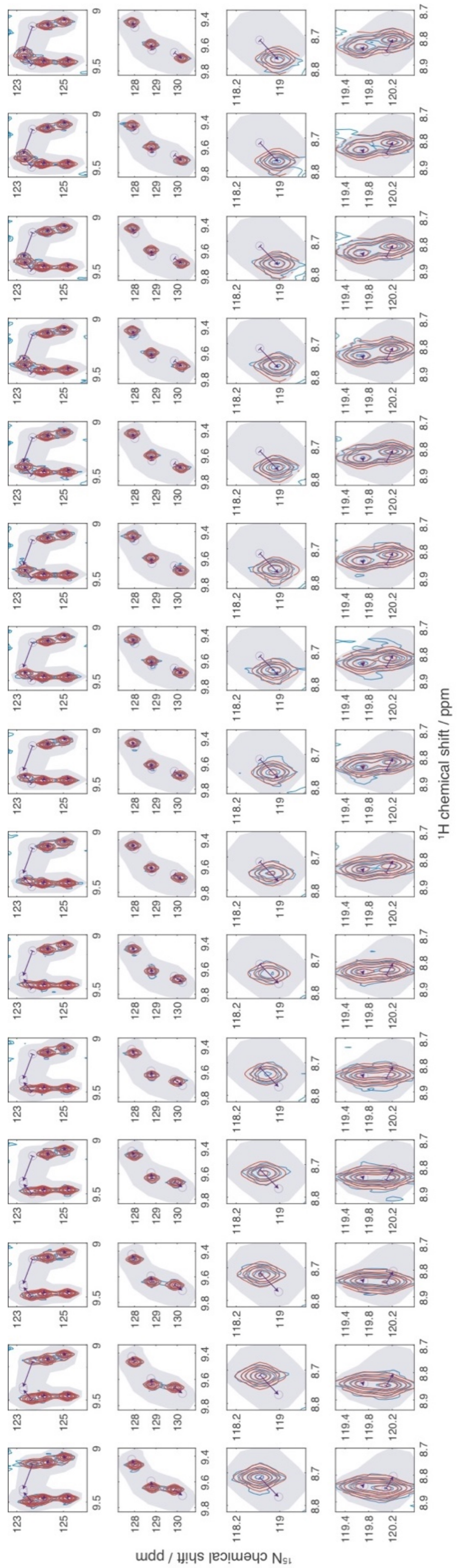

**Fig. S3.** $^{1}$H,$^{15}$N SOFAST-HMQC measurements (blue) and TITAN fits (red) of Hsp90 titration with compound **1** (Fig. 5). Grey shading indicates fitted ROIs.

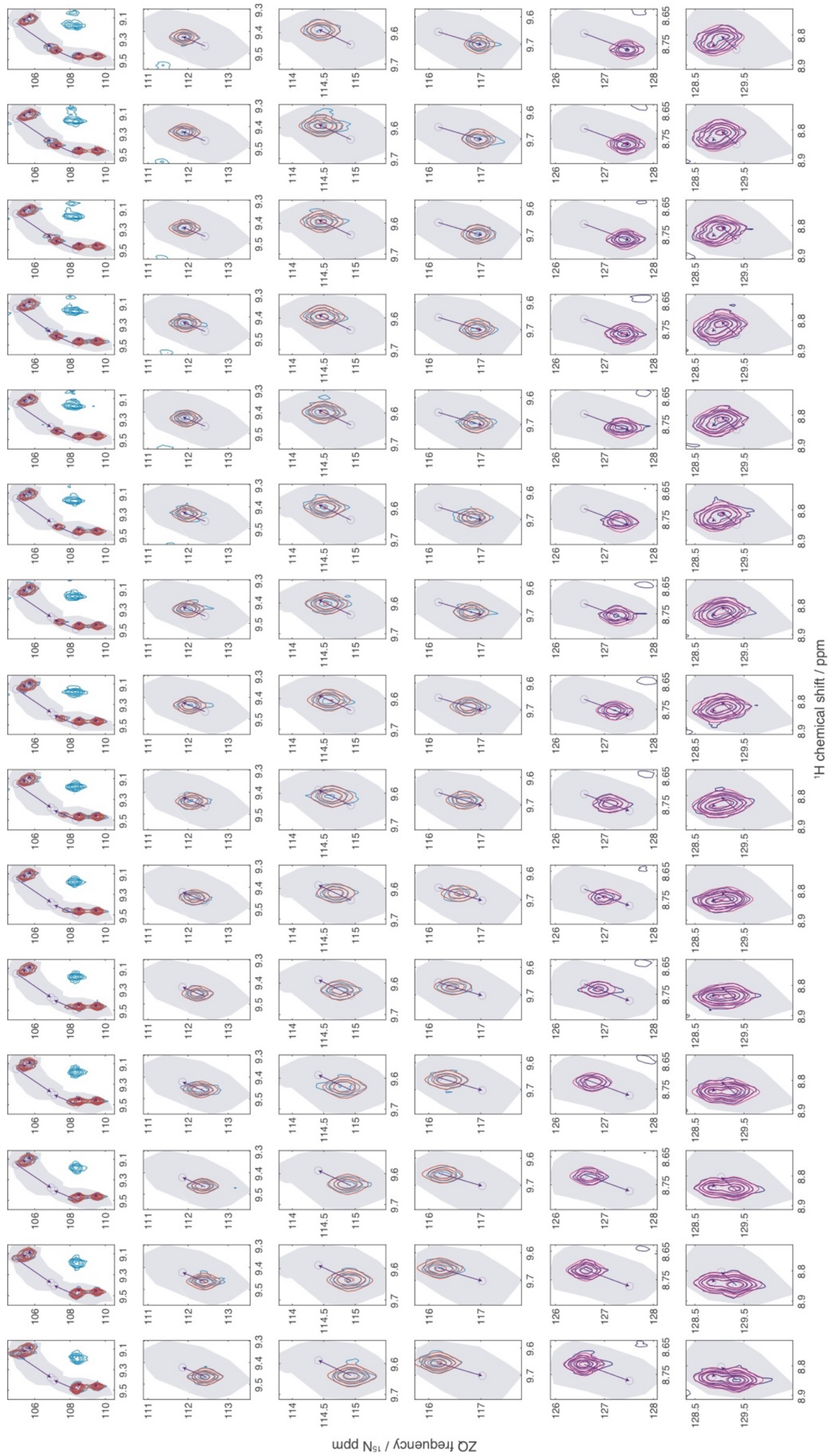

**Fig. S4.** $^{1}H$,$^{15}N$ SOFAST-HZQC measurements (blue/navy) and TITAN fits (red/magenta) of Hsp90 titration with compound **1** (Fig. 5). Grey shading indicates fitted ROIs. Navy/magenta contours indicate negative (folded) crosspeaks.

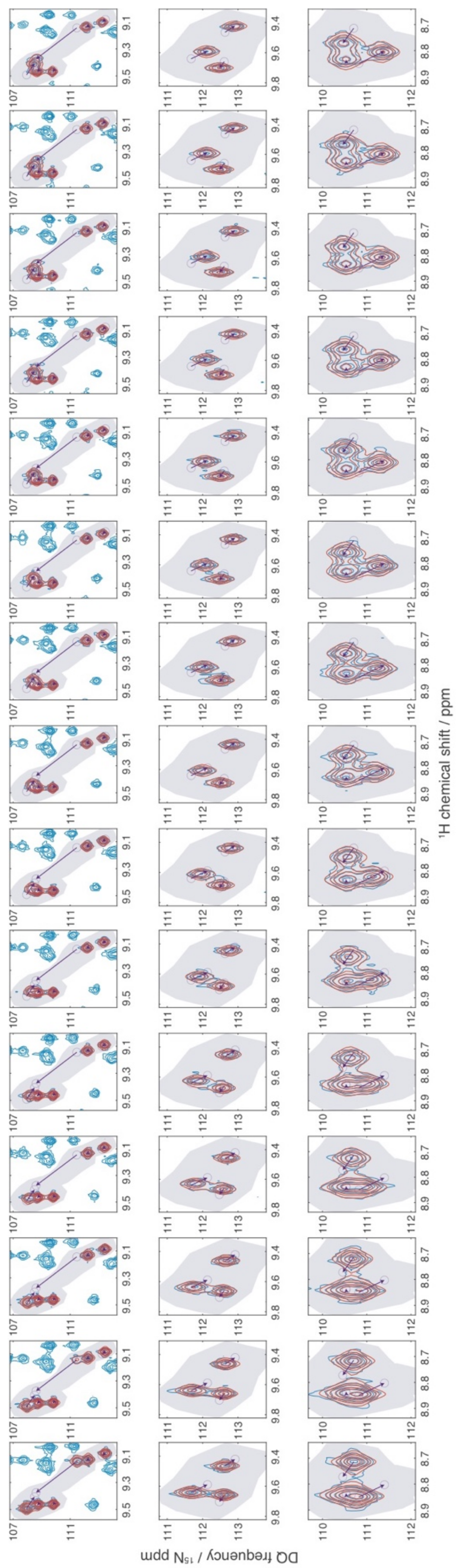

**Fig. S5.** $^{1}$H,$^{15}$N SOFAST-HDQC measurements (blue) and TITAN fits (red) of Hsp90 titration with compound **1** (Fig. 5). Grey shading indicates fitted ROIs.

**Listing S1: SOFAST-H(Z/D)QC pulse program**

```
;SOFAST-H(Z/D)QC
;  Waudby, Ouvry, Davis & Christodoulou (submitted, 2019)
;  for simultaneous collection of SOFAST-HZQC and SOFAST-HDQC spectra
;
;run as pseudo-3D (td1 = 2)
;process using nmrPipe scripts supplied

prosol relations=<triple>

#include <Avance.incl>
#include <Grad.incl>
#include <Delay.incl>

"d11=30m"
"d12=20u"
"d13=4u"
"d21=1s/(cnst4*2)"

"in0=inf2"
"td1=2"
"l0=1"
"acqt0=de"

# ifdef ONE_D
"d0=0.1u"
#else
"d0=in0/2-p21*4/3.1415"
# endif /*ONE_D*/

"DELTA1=d21-p16-d16-p39*cnst39"
"DELTA2=p39*cnst39-de-4u"
"DELTA3=DELTA1-p40*0.5"

# ifdef OFFRES_PRESAT
  "TAU=d1-10m-60u-d12*2-d13"
# else
  "TAU=d1-10m"
# endif /*OFFRES_PRESAT*/

"spoff23=bf1*(cnst19/1000000)-o1"
"spoff24=bf1*(cnst19/1000000)-o1"

aqseq 312

1 ze
  d11 pl26:f3
2 10m do:f3

# ifdef OFFRES_PRESAT
  30u fq=cnst21(bf hz):f1
  d12 pl9:f1
  TAU cw:f1 ph29
  d13 do:f1
  d12 pl1:f1
  30u fq=0:f1
```

```
# else
  TAU
# endif /*OFFRES_PRESAT*/

  d12 pl3:f3
  50u UNBLKGRAD

  ; purge Nz
  (p21 ph1):f3
  p16:gp2
  d16

  ; begin main sequence
  (p39:sp23 ph11):f1
  p16:gp1
  d16

  if "l0 %2 == 1"
     {
  ; 1H 180 before d0
  (lalign (DELTA3 p40:sp24 ph1) (DELTA1 p21 ph12 d0 p21 ph1 DELTA1):f3
)
     }
  else
     {
  ; 1H 180 after d0
  (ralign (p40:sp24 ph1 DELTA3 ) (DELTA1 p21 ph12 d0 p21 ph1 DELTA1):f3
)
     }

  DELTA2
  p16:gp1
  d16 pl26:f3
  4u BLKGRAD

  go=2 ph31 cpd3:f3
  10m do:f3 mc #0 to 2
     F1QF(ip12)
     F2EA(rp12 & iu0, id0)

exit

ph1=0
ph2=0
ph11=0
ph12=0 2
ph31=0 2

;pl3 : f3 channel - power level for pulse (default)
;pl9 : f1 channel - power level for presaturation
;pl26: f3 channel - power level for CPD/BB decoupling (low power)
;sp23: f1 channel - shaped pulse 120 degree (Pc9_4_120.1000)
;sp24: f1 channel - shaped pulse 180 degree (Reburp.1000)
;p16: homospoil/gradient pulse                       [1 msec]
;p21: f3 channel -  90 degree high power pulse
;p39: f1 channel - 120 degree shaped pulse for excitation
;                      Pc9_4_120.1000 (120o)    (2657us at 700 MHz)
```

```
;p40: f1 channel - 180 degree shaped pulse for refocussing
;                       Reburp.1000              (1935us at 700 MHz)
;d0 : incremented delay (2D) = in0/2-p21*4/3.1415
;d1 : relaxation delay
;d11: delay for disk I/O                                [30 msec]
;d12: delay for power switching                         [20 usec]
;d16: delay for homospoil/gradient recovery
;d21 : 1/(2J)NH
;o1 : recommended to place on H(N) = cnst19
;cnst4: = J(NH)
;cnst19: H(N) chemical shift (offset, in ppm)  [8.2 ppm]
;cnst21: frequency (in Hz) for off-resonance presaturation
;cnst39: compensation of chemical shift evolution during p39
;           Pc9_4_120.1000: 0.529
;NS: 2 * n
;DS: 16
;aq: <= 100 msec (with low power cpd, max. 50% duty cycle)
;td1: 2
;td2: number of experiments
;FnMODE[1]: QF
;FnMODE[2]: Echo-AntiEcho
;cpd3: decoupling according to sequence defined by cpdprg3: garp4.p62
;pcpd3: f3 channel - 90 degree pulse for decoupling sequence
;          use pulse of >= 350 usec

;Options:
;  -DOFFRES_PRESATURATION : off-resonance presaturation during d1
(cnst21, pl9)
;  -DONE_D : for 1D measurement with minimal evolution time (td2=1,
td1=2)

;use gradient ratio:    gp 1 : gp 2
;              11 :    7
;
;for z-only gradients:
;gpz1: 11%
;gpz2:  7%
;
;use gradient files:
;gpnam1: SMSQ10.100
;gpnam2: SMSQ10.100
```

**Listing S2: SOFAST-H(Z/D)QC parameter file**

```
##TITLE= Parameter file, TopSpin 3.5 pl 6
##JCAMPDX= 5.0
##DATATYPE= Parameter Values
##NPOINTS= 12   $$ modification sequence number
##ORIGIN= Bruker BioSpin GmbH
##OWNER= waudbyc
$$ 2018-11-03 12:16:10.622 +0000  waudbyc@cl-nmr-spec701
$$ /home/waudbyc/nmr/chris_hzdqc_sensitivity_031118/5/acqus
$$ process /opt/topspin3.5pl6/prog/mod/go4
##$ACQT0= 10
##$AMP= (0..31)
100 100 100 100 100 100 100 100 100 100 100 100 100 100 100 100 100 100
100 100 100 100 100 100 100 100 100 100 100 100 100 100
##$AMPCOIL= (0..19)
0 0 0 0 0 0 0 0 0 0 0 0 0 0 0 0 0 0 0 0
##$ANAVPT= -1
##$AQSEQ= 1
##$AQ_mod= 3
##$AUNM= <au_zg>
##$AUTOPOS= <>
##$BF1= 700.13
##$BF2= 176.047828526
##$BF3= 70.943556786
##$BF4= 700.13
##$BF5= 700.13
##$BF6= 700.13
##$BF7= 700.13
##$BF8= 700.13
##$BWFAC= (0..63)
0 0 0 0 0 0 0 0 0 0 0 0 0 0 0 0 0 0 0 0 0 0 0 0 0 0 0 0 0 0 0 0 0 0 0 0
0 0 0 0 0 0 0 0 0 0 0 0 0 0 0 0 0 0 0 0 0 0 0 0 0 0 0 0
##$BYTORDA= 0
##$CAGPARS= (0..11)
0 0 0 0 0 0 0 0 0 0 0 0
##$CHEMSTR= <none>
##$CNST= (0..63)
1 1 1 1 90 1 1 1 1 1 1 1 1 1 1 1 1 1 1 8.2 1 1 1 1 1 1 1 1 1 1 1 1 1 1
1 1 1 1 1 0.529 1 1 1 1 1 1 1 1 1 1.074 1 1 1 1 8.2 5 1 1 1 1 1 1 1 1
##$CPDPRG= (0..8)
<> <> <> <garp4.p62> <> <> <> <> <>
##$D= (0..63)
0.000155462 0.2 0 0 0 0 0 0 0 0 0 0.03 2e-05 4e-06 0 0 0.0002 0 0 0 0
0.005555556
0 0 0 0 0 0 0 0 0 0 0 0 0 0 0 0 0 0 0 0 0 0 0 0 0 0 0 0 0 0 0 0 0 0 0 0
0 0 0 0 0 0
##$DATE= 1541246423
##$DE= 14.25003
##$DECBNUC= <off>
##$DECIM= 1904
##$DECNUC= <off>
##$DECSTAT= 4
##$DIGMOD= 3
##$DIGTYP= 12
##$DQDMODE= 0
##$DR= 22
```

```
##$DS= 64
##$DSPFIRM= 4
##$DSPFVS= 21
##$DTYPA= 0
##$EXP= <SFHMQCF3GPPH>
##$FCUCHAN= (0..9)
0 2 1 3 0 0 0 0 0 0
##$FL1= 0
##$FL2= 0
##$FL3= 0
##$FL4= 0
##$FN_INDIRECT= (0..7)
0 2 1 0 0 0 0 0
##$FOV= 0
##$FQ1LIST= <>
##$FQ2LIST= <>
##$FQ3LIST= <>
##$FQ4LIST= <>
##$FQ5LIST= <>
##$FQ6LIST= <>
##$FQ7LIST= <>
##$FQ8LIST= <>
##$FRQLO3= 491711
##$FRQLO3N= 2
##$FS= (0..7)
83 83 83 83 83 83 83 83
##$FTLPGN= 0
##$FW= 4032000
##$FnILOOP= 0
##$FnMODE= 0
##$FnTYPE= 0
##$GPNAM= (0..31)
<> <SMSQ10.100> <SMSQ10.100> <> <> <> <> <> <> <> <> <> <> <> <> <> <>
<> <> <> <> <> <> <> <> <> <> <> <> <> <> <>
##$GPX= (0..31)
0 0 0 0 0 0 0 0 0 0 0 0 0 0 0 0 0 0 0 0 0 0 0 0 0 0 0 0 0 0 0 0
##$GPY= (0..31)
0 0 0 0 0 0 0 0 0 0 0 0 0 0 0 0 0 0 0 0 0 0 0 0 0 0 0 0 0 0 0 0
##$GPZ= (0..31)
0 11 7 0 0 0 0 0 0 0 0 0 0 0 0 0 0 0 0 0 0 0 0 0 0 0 0 0 0 0 0 0
##$GRDPROG= <grad_out>
##$GRPDLY= 76
##$HDDUTY= 20
##$HDRATE= 1
##$HGAIN= (0..3)
0 0 0 0
##$HL1= 0
##$HL2= 0
##$HL3= 0
##$HL4= 0
##$HOLDER= 0
##$HPMOD= (0..7)
0 0 0 0 0 0 0 0
##$HPPRGN= 0
##$IN= (0..63)
0.0004026 0 0 0 0 0 0 0 0 0 0 0 0 0 0 0 0 0 0 0 0 0 0 0 0 0 0 0 0 0 0 0
0 0 0 0 0 0 0 0 0 0 0 0 0 0 0 0 0 0 0 0 0 0 0 0 0 0 0 0 0 0 0 0
```

##$INF= (0..7)
0 142.8 402.6 0 0 0 0 0
##$INP= (0..63)
0 0 0 0 0 0 0 0 0 0 0 0 0 0 0 0 0 0 0 0 0 0 0 0 0 0 0 0 0 0 0 0 0 0 0 0
0 0 0 0 0 0 0 0 0 0 0 0 0 0 0 0 0 0 0 0 0 0 0 0 0 0 0 0
##$INSTRUM= <spect>
##$INTEGFAC= (0..63)
0 0 0 0 0 0 0 0 0 0 0 0 0 0 0 0 0 0 0 0 0 0 0 0 0 0 0 0 0 0 0 0 0 0 0 0
0 0 0 0 0 0 0 0 0 0 0 0 0 0 0 0 0 0 0 0 0 0 0 0 0 0 0 0
##$L= (0..31)
1 1 1 1 1 1 1 1 1 1 1 1 1 1 1 1 1 1 1 1 1 1 1 1 1 1 1 1 1 1 1 1
##$LFILTER= 100
##$LGAIN= -5
##$LINPSTP= 0
##$LOCKED= yes
##$LOCKFLD= -997
##$LOCKGN= 116.600120544434
##$LOCKPOW= -16
##$LOCKPPM= 4.69999980926514
##$LOCNUC= <2H>
##$LOCPHAS= 239.2514
##$LOCSHFT= yes
##$LOCSW= 0
##$LTIME= 0.349999994039536
##$MASR= 4200
##$MASRLST= <masrlst>
##$MULEXPNO= (0..15)
0 0 0 0 0 0 0 0 0 0 0 0 0 0 0 0
##$NBL= 1
##$NC= -6
##$NLOGCH= 1
##$NOVFLW= 0
##$NS= 2
##$NUC1= <1H>
##$NUC2= <13C>
##$NUC3= <15N>
##$NUC4= <off>
##$NUC5= <off>
##$NUC6= <off>
##$NUC7= <off>
##$NUC8= <off>
##$NUCLEUS= <off>
##$NUSLIST= <automatic>
##$NusAMOUNT= 50
##$NusFPNZ= no
##$NusJSP= 0
##$NusSEED= 54321
##$NusSPTYPE= 0
##$NusT2= 1
##$NusTD= 0
##$O1= 3289
##$O2= 17780.831
##$O3= 8300.396143962
##$O4= 3290.611
##$O5= 3290.611
##$O6= 3290.611
##$O7= 3290.611

```
##$O8= 3290.611
##$OVERFLW= 0
##$P= (0..63)
7.6 10.68 15.2 11.2 22.4 16.5 25 50 500 25 50 1000 2000 300 231 1602
1000
2500 100000 600 2000 36 72 1286 900 129 65 7.6 0 4000 160 694 0 0 0 137
0 0 0 2657 1935 1886 1200 1457 171 78 52 1800 2829 85714 5143 553 1200
0 0 1114 500 1371 0 0 343 110 350 1500
##$PACOIL= (0..15)
0 0 0 0 0 0 0 0 0 0 0 0 0 0 0 0
##$PAPS= 0
##$PARMODE= 2
##$PCPD= (0..9)
0 65 55 220 0 0 0 0 0 0
##$PEXSEL= (0..9)
1 1 1 1 1 1 1 1 1 1
##$PHCOR= (0..31)
0 0 0 0 0 0 0 0 0 0 0 0 0 0 0 0 0 0 0 0 0 0 0 0 0 0 0 0 0 0 0 0
##$PHLIST= <>
##$PHP= 1
##$PH_ref= 0
##$PL= (0..63)
120 120 120 120 120 120 120 120 120 120 120 120 120 120 120 120 120 120
120 120 120 120 120 120 120 120 120 120 120 120 120 120 120 120 120 120
120 120 120 120 120 120 120 120 120 120 120 120 120 120 120 120 120 120
120 120 120 120 120 120 120 120 120 120
##$PLSTEP= 0.1
##$PLSTRT= -6
##$PLW= (0..63)
0 7 113 98 0 0 0 0 0 1.6173e-05 0.64691 0.063175 4.6859 0 1.1715 22.68
2.6241 0 7 0.095697 2.3298 0.14953 5.2421 19.845 0.0252 7.5153 1.0351
7.6322
8.1094 0.12048 4.6859 18.743 6.4691e-07 0 0 0 10.497 0 0 0 0 0 0 0 0 0
0 0 0 0 0 0 0 0 0 0 0 0 0 0 0 0 0 0
##$PLWMAX= (0..7)
84.2 385 482.3 0 0 0 0 0
##$PQPHASE= 0
##$PQSCALE= 1
##$PR= 1
##$PRECHAN= (0..15)
-1 6 0 1 4 5 -1 -1 -1 -1 -1 -1 -1 -1 -1 -1
##$PRGAIN= 0
##$PROBHD= <Z142131_0001 (CP QCI 700S4 H&F/P/C-N-D-05 Z)>
##$PULPROG= <sfhzdqcf3.cw>
##$PW= 0
##$PYNM= <>
##$ProjAngle= 0
##$QNP= 0
##$RD= 0
##$RECCHAN= (0..15)
0 2 1 0 0 0 0 0 0 0 0 0 0 0 0 0
##$RECPH= 0
##$RECPRE= (0..15)
-1 6 0 -1 -1 -1 -1 -1 -1 -1 -1 -1 -1 -1 -1 -1
##$RECPRFX= (0..15)
1 0 0 0 0 0 1 0 0 0 0 0 0 0 0 0
##$RECSEL= (0..15)
```

```
0 1 2 0 0 0 0 0 0 0 0 0 0 0 0 0
##$RG= 203
##$RO= 0
##$RSEL= (0..15)
0 1 2 4 0 0 0 0 0 0 0 0 0 0 0 0
##$S= (0..7)
83 83 83 83 83 83 83 83
##$SELREC= (0..9)
0 0 0 0 0 0 0 0 0 0
##$SFO1= 700.133289
##$SFO2= 176.065609357
##$SFO3= 70.951857182144
##$SFO4= 700.133290611
##$SFO5= 700.133290611
##$SFO6= 700.133290611
##$SFO7= 700.133290611
##$SFO8= 700.133290611
##$SOLVENT= <H2O+D2O>
##$SOLVOLD= <off>
##$SP= (0..63)
120 120 120 120 120 120 120 120 120 120 120 120 120 120 120 120 120 120
120 120 120 120 120 120 120 120 120 120 120 120 120 120 120 120 120 120
120 120 120 120 120 120 120 120 120 120 120 120 120 120 120 120 120 120
120 120 120 120 120 120 120 120 120 120
##$SPECTR= 0
##$SPINCNT= 0
##$SPNAM= (0..63)
<> <Sinc1.1000> <Q5_sebop.1> <Q3_surbop.1> <Q5_sebop.1> <Q3_surbop.1>
<Q5tr_sebop.1>
<Q3_surbop.1> <Q5tr_sebop.1> <Q3_surbop.1> <Q5.1000> <Sinc1.1000>
<Q5tr.1000>
<Crp80,0.5,20.1> <Crp42,1.5,20.2> <Q3_surbop.1> <Q3_surbop.1>
<Q3_surbop.1>
<Crp60_xfilt.2> <Iburp2.1000> <Q3_surbop.1> <Reburp.1000>
<Pc9_4_90.1000>
<Pc9_4_120.1000> <Reburp.1000> <Pc9_4_90.1000> <Reburp.1000>
<Pc9_4_90.1000>
<Eburp2.1000> <Eburp2tr.1000> <Bip720,50,20.1> <Crp42,1.5,20.2>
<Gaus1_180i.1000>
<Q3_surbop.1> <Q3.1000> <Reburp.1000> <0.0> <0.0> <Iburp2.1000>
<Bip720,50,20.1>
<Reburp.1000> <Bip720,100,10.1> <> <> <> <> <> <> <> <> <> <> <> <> <>
<> <> <> <> <> <> <> <> <>
##$SPOAL= (0..63)
0.5 0.5 1 0.5 1 0.5 0 0.5 0 0.5 1 0.5 0 0.5 0.5 0.5 0.5 0.5 0.5 0.5 0.5
0.5 1 1 0.5 1 0.5 0 1 0 0.5 0.5 0.5 0.5 0.5 0.5 0.5 0.5 0.5 0.5 0.5 0.5
0.5 0.5 0.5 0.5 0.5 0.5 0.5 0.5 0.5 0.5 0.5 0.5 0.5 0.5 0.5 0.5 0.5 0.5
0.5 0.5 0.5 0.5
##$SPOFFS= (0..63)
0 0 0 0 0 0 0 0 0 0 0 0 0 0 0 0 0 0 0 0 0 0 0 2452.066 2452.066 0 0 0 0
0 0 0 0 0 0 0 0 0 0 0 0 0 0 0 0 0 0 0 0 0 0 0 0 0 0 0 0 0 0 0 0 0 0 0
##$SPPEX= (0..63)
0 0 0 0 0 0 0 0 0 0 0 0 0 0 0 0 0 0 0 0 0 0 0 0 0 0 0 0 0 0 0 0 0 0 0 0
0 0 0 0 0 0 0 0 0 0 0 0 0 0 0 0 0 0 0 0 0 0 0 0 0 0 0 0
##$SPW= (0..63)
0 0.0011658 70.311 73.196 70.311 73.196 70.311 73.196 70.311 4.822
2.8846
```

```
7.2866e-05 2.8846 28.877 12.128 8.1094 4.822 4.822 7.2996 1.7278 73.196
1.1122 0.0009782501 0.01285287 0.1339677 0.0072744 0.17632 0.0072744
0.051136
0.051136 0.88494 3.032 1.2995e-06 4.822 15.375 0.83027 0 0 0.12867
32.514
42.433 48.334 0 0 0 0 0 0 0 0 0 0 0 0 0 0 0 0 0 0 0 0 0 0
##$SUBNAM= (0..9)
<> <> <> <> <> <> <> <> <> <>
##$SW= 15.0031456091389
##$SWIBOX= (0..19)
0 1 2 0 5 0 0 0 0 0 0 0 0 0 0 0 0 0 0 0
##$SW_h= 10504.2016806723
##$SWfinal= 0
##$SigLockShift= 0
##$TD= 4096
##$TD0= 1
##$TD_INDIRECT= (0..7)
0 2 512 0 0 0 0 0
##$TDav= 1
##$TE= 282.9984
##$TE1= 283.9297
##$TE2= 0
##$TE3= 0
##$TE4= 0
##$TEG= 300
##$TE_MAGNET= 0
##$TE_PIDX= 1
##$TE_STAB= (0..9)
2 2 0 0 0 0 0 0 0 0
##$TL= (0..7)
120 120 120 120 120 120 120 120
##$TOTROT= (0..63)
0 0 0 0 0 0 0 0 0 0 0 0 0 0 0 0 0 0 0 0 0 0 0 0 0 0 0 0 0 0 0 0 0 0 0 0
0 0 0 0 0 0 0 0 0 0 0 0 0 0 0 0 0 0 0 0 0 0 0 0 0 0 0 0
##$TUBE_TYPE= <>
##$USERA1= <>
##$USERA2= <>
##$USERA3= <>
##$USERA4= <>
##$USERA5= <>
##$V9= 5
##$VALIDCODE= -1
##$VALIST= <>
##$VCLIST= <>
##$VDLIST= <>
##$VPLIST= <>
##$VTLIST= <>
##$WBST= 1024
##$WBSW= 20
##$XGAIN= (0..3)
0 0 0 0
##$XL= 0
##$YL= 0
##$YMAX_a= 26525
##$YMIN_a= -28758
##$ZGOPTNS= <>
##$ZL1= 120
```

```
##$ZL2= 120
##$ZL3= 120
##$ZL4= 120
##END=
```

**Listing S3: SOFAST-H(Z/D)QC nmrPipe processing template**

```
#!/bin/csh
#
# Requires shufZQ.m and shufDQ.m
#
# yN = 2 * number of t1 points (2*td2)
# yT = number of t1 points (td2)

bruk2pipe -in ./ser \
  -bad 0.0 -ext -aswap -AMX -decim 1904 -dspfvs 21 -grpdly 76  \
  -xN              4096  -yN              1024  \
  -xT              2048  -yT               512  \
  -xMODE            DQD  -yMODE        Complex  \
  -xSW        10504.202  -ySW         2483.855  \
  -xOBS         700.133  -yOBS          70.952  \
  -xCAR           4.916  -yCAR         117.225  \
  -xLAB              HN  -yLAB             15N  \
  -ndim               2  -aq2D         Complex  \
  -out ./test.fid -verb -ov

nmrPipe -in test.fid \
| nmrPipe  -fn MAC -all -noRd -noWr -macro shufZQ.m \
| nmrPipe  -fn SOL \
| nmrPipe  -fn SP -off 0.5 -end 1.00 -pow 1 -c 1.0    \
| nmrPipe  -fn ZF -zf 2                               \
| nmrPipe  -fn FT -auto                               \
| nmrPipe  -fn PS -p0 220.00 -p1 0.00 -di -verb         \
| nmrPipe  -fn EXT -x1 7ppm -xn 9ppm -sw \
| nmrPipe  -fn TP                                     \
| nmrPipe  -fn LP -fb \
| nmrPipe  -fn SP -off 0.5 -end 1.00 -pow 1 -c 1.0    \
| nmrPipe  -fn ZF -auto                               \
| nmrPipe  -fn FT -neg                               \
| nmrPipe  -fn PS -p0 -115.00 -p1 180.00 -di -verb         \
   -ov -out hzqc.ft2

nmrPipe -in test.fid \
| nmrPipe  -fn MAC -all -noRd -noWr -macro shufDQ.m \
| nmrPipe  -fn SOL \
| nmrPipe  -fn SP -off 0.5 -end 1.00 -pow 1 -c 1.0    \
| nmrPipe  -fn ZF -zf 2                               \
| nmrPipe  -fn FT -auto                               \
| nmrPipe  -fn PS -p0 -148.00 -p1 0.00 -di -verb         \
| nmrPipe  -fn EXT -x1 7ppm -xn 9ppm -sw \
| nmrPipe  -fn TP                                     \
| nmrPipe  -fn LP -fb \
| nmrPipe  -fn SP -off 0.5 -end 1.00 -pow 1 -c 1.0    \
| nmrPipe  -fn ZF -auto                               \
| nmrPipe  -fn FT -neg                               \
| nmrPipe  -fn PS -p0 -65.00 -p1 180.00 -di -verb         \
   -ov -out hdqc.ft2
```

**Listing S4: shufZQ.m (nmrPipe processing macro)**

```
/***/
/* shufZQ.M: shuffling SOFAST-H(Z/D)QC input to extract ZQ spectrum
/*
/* Example: | nmrPipe -fn MAC -all -noRd -noWr -macro shufZQ.M \
/***/

/***/
/* Initialization steps:
/*  ARGS: extract summation or difference mode.
/*  INIT: update header to reflect reduced number of 1D vectors.
/***/

if (sliceCode == CODE_ARGS)
   {
   };

if (sliceCode == CODE_INIT)
   {
    tdSize   = getParm( fdata, NDTDSIZE, CUR_YDIM );
    apodSize = getParm( fdata, NDAPOD,   CUR_YDIM );

    (void) setParm( fdata, NDSIZE,   integer( specnum/2 ),  CUR_YDIM );
    (void) setParm( fdata, NDTDSIZE, integer( tdSize/2 ),   CUR_YDIM );
    (void) setParm( fdata, NDAPOD,   integer( apodSize/2 ), CUR_YDIM );
   };

if (sliceCode < 1)
   {
    exit( 0 );
   };

/***/
/* Process only every group of 4 complex 1D slices:
/***/

if (sliceCode % 4)
   {
    exit( 0 );
   };

float vA[size],  vB[size],  vC[size],  vD[size],  vE[size],  vF[size],
vG[size],  vH[size];
float vAA[size], vBB[size], vCC[size], vDD[size];

(void) dReadB( inUnit, vA, wordLen*size );
(void) dReadB( inUnit, vB, wordLen*size );
(void) dReadB( inUnit, vC, wordLen*size );
(void) dReadB( inUnit, vD, wordLen*size );
(void) dReadB( inUnit, vE, wordLen*size );
(void) dReadB( inUnit, vF, wordLen*size );
(void) dReadB( inUnit, vG, wordLen*size );
(void) dReadB( inUnit, vH, wordLen*size );

/***/
/* Desired output:
```

```
/*    cos component = (fid 1)  + i(fid 2)  + (fid 3)  - i(fid 4)
/*                  = (A + iB) + i(C + iD) + (E + iF) - i(G + iH)
/*
/*    sin component = i(fid 1)  - (fid 2)  - i(fid 3)  - (fid 4)
/*                  = i(A + iB) - (C + iD) - i(E + iF) - (G + iH)
/*
/* AA = (t1 = 1 real, t2 = real) = A + (-D) + E - (-H)
/* BB = (t1 = 1 real, t2 = imag) = B + C + F - G
/*
/* CC = (t1 = 2 imag, t2 = real) = (-B) - C - (-F) - G
/* DD = (t1 = 2 imag, t2 = imag) = A - D - E - H
/***/

(void) vvCopy( vAA, vA, size );
(void) vvSub( vAA, vD, size );
(void) vvAdd( vAA, vE, size );
(void) vvAdd( vAA, vH, size );

(void) vvCopy( vBB, vB, size );
(void) vvAdd( vBB, vC, size );
(void) vvAdd( vBB, vF, size );
(void) vvSub( vBB, vG, size );

(void) vvCopy( vCC, vB, size );
(void) vNeg( vCC, size );
(void) vvSub( vCC, vC, size );
(void) vvAdd( vCC, vF, size );
(void) vvSub( vCC, vG, size );

(void) vvCopy( vDD, vA, size );
(void) vvSub( vDD, vD, size );
(void) vvSub( vDD, vE, size );
(void) vvSub( vDD, vH, size );

(void) dWrite( outUnit, vAA, wordLen*size );
(void) dWrite( outUnit, vBB, wordLen*size );
(void) dWrite( outUnit, vCC, wordLen*size );
(void) dWrite( outUnit, vDD, wordLen*size );
```

**Listing S5: shufDQ.m  (nmrPipe processing macro)**

```
/***/
/* shufDQ.M: shuffling SOFAST-H(Z/D)QC input to extract DQ spectrum
/*
/* Example: | nmrPipe -fn MAC -all -noRd -noWr -macro shufDQ.M \
/***/

/***/
/* Initialization steps:
/*  ARGS: extract summation or difference mode.
/*  INIT: update header to reflect reduced number of 1D vectors.
/***/

if (sliceCode == CODE_ARGS)
   {
   };

if (sliceCode == CODE_INIT)
   {
    tdSize   = getParm( fdata, NDTDSIZE, CUR_YDIM );
    apodSize = getParm( fdata, NDAPOD,   CUR_YDIM );

    (void) setParm( fdata, NDSIZE,   integer( specnum/2 ),  CUR_YDIM );
    (void) setParm( fdata, NDTDSIZE, integer( tdSize/2 ),   CUR_YDIM );
    (void) setParm( fdata, NDAPOD,   integer( apodSize/2 ), CUR_YDIM );
   };

if (sliceCode < 1)
   {
    exit( 0 );
   };

/***/
/* Process only every group of 4 complex 1D slices:
/***/

if (sliceCode % 4)
   {
    exit( 0 );
   };

float vA[size],  vB[size],  vC[size],  vD[size],  vE[size],  vF[size],
vG[size],  vH[size];
float vAA[size], vBB[size], vCC[size], vDD[size];

(void) dReadB( inUnit, vA, wordLen*size );
(void) dReadB( inUnit, vB, wordLen*size );
(void) dReadB( inUnit, vC, wordLen*size );
(void) dReadB( inUnit, vD, wordLen*size );
(void) dReadB( inUnit, vE, wordLen*size );
(void) dReadB( inUnit, vF, wordLen*size );
(void) dReadB( inUnit, vG, wordLen*size );
(void) dReadB( inUnit, vH, wordLen*size );

/***/
/* Desired output:
```

```
/*    cos component = (fid 1)  - i(fid 2)  + (fid 3)  + i(fid 4)
/*                  = (A + iB) - i(C + iD) + (E + iF) + i(G + iH)
/*
/*    sin component = -i(fid 1)  - (fid 2)  + i(fid 3)  - (fid 4)
/*                  = -i(A + iB) - (C + iD) + i(E + iF) - (G + iH)
/*
/* AA = (t1 = 1 real, t2 = real) = A + D + E - H
/* BB = (t1 = 1 real, t2 = imag) = B - C + F + G
/*
/* CC = (t1 = 2 imag, t2 = real) = B - C - F - G
/* DD = (t1 = 2 imag, t2 = imag) = (-A) - D + E - H
/***/

(void) vvCopy( vAA, vA, size );
(void) vvAdd( vAA, vD, size );
(void) vvAdd( vAA, vE, size );
(void) vvSub( vAA, vH, size );

(void) vvCopy( vBB, vB, size );
(void) vvSub( vBB, vC, size );
(void) vvAdd( vBB, vF, size );
(void) vvAdd( vBB, vG, size );

(void) vvCopy( vCC, vB, size );
(void) vvSub( vCC, vC, size );
(void) vvSub( vCC, vF, size );
(void) vvSub( vCC, vG, size );

(void) vvCopy( vDD, vA, size );
(void) vNeg( vDD, size );
(void) vvSub( vDD, vD, size );
(void) vvAdd( vDD, vE, size );
(void) vvSub( vDD, vH, size );

(void) dWrite( outUnit, vAA, wordLen*size );
(void) dWrite( outUnit, vBB, wordLen*size );
(void) dWrite( outUnit, vCC, wordLen*size );
(void) dWrite( outUnit, vDD, wordLen*size );
```

**Listing S6: BEST-ZQ/DQ-TROSY pulse program**

```
;BEST-TROSY-H(Z/D)QC
;  Waudby, Ouvry, Davis & Christodoulou (submitted, 2019)
;
;options:
; -DLABEL_CN = 13C decoupling
; -DDQ = HDQC (otherwise runs HZQC)
; -DONE_D = first-row
; -DOFFRES_PRESAT = presat, pl9 on cnst21 (Hz bf)

prosol relations=<triple>

#include <Avance.incl>
#include <Grad.incl>
#include <Delay.incl>

"d11=30m"
"d12=20u"
"d13=4u"
"d21=1s/(cnst4*4)"

"p22=p21*2"

"in0=inf1"
# ifdef ONE_D
"d0=2u"
#else
"d0=in0/2-p21*4/3.1415"
# endif /*ONE_D*/

"d2=p39-p39*cnst39-0.3633*p21"
"d3=0.5*p40-0.3633*p21"
"DELTA1=d21-p39*cnst39-p40*0.5-p16-d16-4u"
"DELTA2=d21-0.3633*p21-p16-d16-4u-0.5*p40"
"DELTA3=d21-p40-p16-d16-4u"
"DELTA4=d21-0.5*p40-p16-d16-4u-p21-de"
"acqt0=de"

#ifdef LABEL_CN
"d10=DELTA3+d3+p21+d0*0.5-p8*0.5"
"d9=DELTA3+d3+p21+d0*0.5-p8*0.5"
"in10=in0*0.5"
"in9=in0*0.5"
#endif

# ifdef OFFRES_PRESAT
  "TAU=d1-d11-60u-d12*2-d13-d12-50u-p21-2*p16-2*d16-12u"
# else
  "TAU=d1-d11-d12-50u-p21-2*p16-2*d16-12u"
# endif /*OFFRES_PRESAT*/


;"spoff23=bf1*(cnst19/1000000)-o1"
;"spoff24=bf1*(cnst19/1000000)-o1"
"spoff23=0" ; for amides on-resonance (recommended)
"spoff24=0"
```

```
"l0=1"  ; loop counter for shifting 1H 180 pulse between echo/anti-
echoes

1 ze
  d11
2 d11

  4u UNBLKGRAD
  p16:gp3
  d16
  4u BLKGRAD

# ifdef OFFRES_PRESAT
  30u fq=cnst21(bf hz):f1
  d12 pl9:f1
  TAU cw:f1 ph1
  d13 do:f1
  d12 pl1:f1
  30u fq=0:f1
# else
  TAU
# endif /*OFFRES_PRESAT*/

3 d12 pl3:f3
  50u UNBLKGRAD

  ; purge Nz
  (p21 ph1):f3
  4u
  p16:gp0
  d16

  ; begin main sequence
  if "l0 %2 == 1"
     {
  (p39:sp23 ph10) (d2 p21 ph11):f3
     }
  else
     {
  (p39:sp23 ph10) (d2 p21 ph21):f3
     }

  DELTA1
  4u
  p16:gp1
  d16
  (center (p40:sp24 ph1) (p22 ph12):f3 )
  4u
  p16:gp1
  d16

  if "l0 %2 == 1"
     {
#ifdef LABEL_CN
  (ralign (p40:sp24 ph16 DELTA3) (DELTA2 p21 ph13 d0 p21 ph1 d3
DELTA3):f3 (p8:sp13 ph1 d10):f2 )
```

```
#else
  (ralign (p40:sp24 ph16) (DELTA2 p21 ph13 d0 p21 ph1 d3):f3 )
  DELTA3
#endif /*LABEL_CN*/
     }
  else
     {
#ifdef LABEL_CN
  (DELTA3 p40:sp24 ph16) (DELTA3 d3 p21 ph23 d0 p21 ph1 DELTA2):f3 (d9
p8:sp13 ph1):f2
#else
  DELTA3
  (p40:sp24 ph16) (d3 p21 ph23 d0 p21 ph1 DELTA2):f3
#endif /*LABEL_CN*/
     }
  4u
  p16:gp2
  d16
  (center (p40:sp24 ph1) (p22 ph1):f3 )
  4u
  p16:gp2
  d16
  DELTA4 BLKGRAD
  (p21 ph14):f3

  go=2 ph31
#ifdef LABEL_CN
  d11 mc #0 to 2
     F1EA(iu0 & ip13*2 & ip14*2, id0 & id10 & id9 & ip10*2 & ip31*2)
#else
  d11 mc #0 to 2
     F1EA(iu0 & ip13*2 & ip14*2, id0 & ip10*2 & ip31*2)
#endif /*LABEL_CN*/

exit

ph1=0
ph10=0
ph11=2 0 3 1
ph21=2 0 1 3
ph12=0
#ifdef DQ
ph13=1 3 2 0
ph23=1 3 0 2
ph14=3
#else /* ZQ */
ph13=1 3 0 2
ph23=1 3 2 0
ph14=1
#endif
ph16=0 0 0 0 1 1 1 1 2 2 2 2 3 3 3 3
ph31=0 2 3 1 2 0 1 3

;pl3 : f3 channel - power level for pulse (default)
;pl9 : f1 channel - power level for presaturation
;pl26: f3 channel - power level for CPD/BB decoupling (low power)
;sp23: f1 channel - shaped pulse 90 degree (Pc9_4_90.1000)
```

```
;sp24: f1 channel - shaped pulse 180 degree (Reburp.1000)
;p16: homospoil/gradient pulse                       [1 msec]
;p21: f3 channel -  90 degree high power pulse
;p39: f1 channel - 90 degree shaped pulse for excitation
;                      Pc9_4_120.1000 (120o)    (1958us at 950 MHz)
;p40: f1 channel - 180 degree shaped pulse for refocussing
;                      Reburp.1000              (1432us at 950 MHz)
;d0 : incremented delay (2D) = in0/2-p21*4/3.1415
;d1 : relaxation delay
;d11: delay for disk I/O                             [30 msec]
;d12: delay for power switching                      [20 usec]
;d16: delay for homospoil/gradient recovery
;d21 : 1/(4J)NH
;cnst4: = J(NH)
;cnst19: H(N) chemical shift (offset, in ppm)  [8.2 ppm]
;cnst21: frequency (in Hz) for off-resonance presaturation
;cnst39: compensation of chemical shift evolution during p39
;           Pc9_4_90.1000: 0.514
;NS: 4 * n
;DS: 16
;td1: number of experiments
;FnMODE: Echo-AntiEcho

;use gradient ratio:    gp 0 : gp 1 : gp 2
;           -16 :  11 :    7

;for z-only gradients:
;gpz0: -16%
;gpz1: 11%
;gpz2:  7%
;gpz3: -23%

;use gradient files:
;gpnam0: SMSQ10.100
;gpnam1: SMSQ10.100
;gpnam2: SMSQ10.100
;gpnam3: SMSQ10.100

;Processing
;PHC0(F1): 90
;PHC1(F1): -180
;FCOR(F1): 1
```

**Listing S7: Parameter file for BEST-ZQ-TROSY**

```
##TITLE= Parameter file, TopSpin 3.5 pl 6
##JCAMPDX= 5.0
##DATATYPE= Parameter Values
##NPOINTS= 11   $$ modification sequence number
##ORIGIN= Bruker BioSpin GmbH
##OWNER= waudbyc
$$ 2019-08-13 19:30:58.188 +0100  waudbyc@cl-nmr-spec950
$$ /home/waudbyc/nmr/chris_ubq_bzqtrosy_130819/13/acqus
$$ process /opt/topspin3.5pl6/prog/mod/go4
##$ACQT0= 10
##$AMP= (0..31)
100 100 100 100 100 100 100 100 100 100 100 100 100 100 100 100 100 100
100 100 100 100 100 100 100 100 100 100 100 100 100 100
##$AMPCOIL= (0..19)
0 0 0 0 0 0 0 0 0 0 0 0 0 0 0 0 0 0 0 0
##$ANAVPT= -1
##$AQSEQ= 0
##$AQ_mod= 3
##$AUNM= <au_zg>
##$AUTOPOS= <>
##$BF1= 950.45
##$BF2= 238.990843
##$BF3= 96.308262
##$BF4= 950.45
##$BF5= 950.45
##$BF6= 950.45
##$BF7= 950.45
##$BF8= 950.45
##$BWFAC= (0..63)
0 0 0 0 0 0 0 0 0 0 0 0 0 0 0 0 0 0 0 0 0 0 0 0 0 0 0 0 0 0 0 0 0 0 0 0
0 0 0 0 0 0 0 0 0 0 0 0 0 0 0 0 0 0 0 0 0 0 0 0 0 0 0 0
##$BYTORDA= 0
##$CAGPARS= (0..11)
0 0 0 0 0 0 0 0 0 0 0 0
##$CHEMSTR= <none>
##$CNST= (0..63)
1 1 1 1 90 1 1 1 1 1 1 1 1 1 1 1 1 1 1 8.2 1 1 1 1 1 1 1 1 1 1 1 1 1 1
1 1 1 1 1 0.529 1 1 1 1 1 1 1 1 1 1.074 1 1 1 1 8.3 5 1 1 1 1 1 1 1 1
##$CPDPRG= (0..8)
<> <> <> <garp4.p62> <> <> <> <> <>
##$D= (0..63)
0.0001214551 0.1 0.0009105501 0.0007042344 0 0 0 0 0 0 0 0.03 2e-05 4e-
06
0 0 0.0002 0 0 0 0 0.002777778 0 0 0 0 0 0 0 0 0 0 0 0 0 0 0 0 0 0 0 0
0 0 0 0 0 0 0 0 0 0 0 0 0 0 0 0 0 0 0 0 0 0
##$DATE= 1565718954
##$DE= 12.02504
##$DECBNUC= <off>
##$DECIM= 1312
##$DECNUC= <off>
##$DECSTAT= 4
##$DIGMOD= 3
##$DIGTYP= 12
##$DQDMODE= 0
##$DR= 22
```

```
##$DS= 128
##$DSPFIRM= 4
##$DSPFVS= 21
##$DTYPA= 0
##$EXP= <SFHMQCF3GPPH>
##$FCUCHAN= (0..9)
0 2 1 4 0 0 0 0 0 0
##$FL1= 0
##$FL2= 0
##$FL3= 0
##$FL4= 0
##$FN_INDIRECT= (0..7)
0 2 0 0 0 0 0 0
##$FOV= 0
##$FQ1LIST= <>
##$FQ2LIST= <>
##$FQ3LIST= <>
##$FQ4LIST= <>
##$FQ5LIST= <>
##$FQ6LIST= <>
##$FQ7LIST= <>
##$FQ8LIST= <>
##$FRQLO3= 479706.309999943
##$FRQLO3N= 2
##$FS= (0..7)
83 83 83 83 83 83 83 83
##$FTLPGN= 0
##$FW= 4032000
##$FnILOOP= 0
##$FnMODE= 0
##$FnTYPE= 0
##$GPNAM= (0..31)
<SMSQ10.100> <SMSQ10.100> <SMSQ10.100> <SMSQ10.100> <> <> <> <> <> <>
<>
<> <> <> <> <> <> <> <> <> <> <> <> <> <> <> <> <> <> <> <> <>
##$GPX= (0..31)
0 0 0 0 0 0 0 0 0 0 0 0 0 0 0 0 0 0 0 0 0 0 0 0 0 0 0 0 0 0 0 0
##$GPY= (0..31)
0 0 0 0 0 0 0 0 0 0 0 0 0 0 0 0 0 0 0 0 0 0 0 0 0 0 0 0 0 0 0 0
##$GPZ= (0..31)
-16 11 7 -23 0 0 0 0 0 0 0 0 0 0 0 0 0 0 0 0 0 0 0 0 0 0 0 0 0 0 0 0
##$GRDPROG= <grad_out>
##$GRPDLY= 76
##$HDDUTY= 20
##$HDRATE= 1
##$HGAIN= (0..3)
0 0 0 0
##$HL1= 0
##$HL2= 0
##$HL3= 0
##$HL4= 0
##$HOLDER= 0
##$HPMOD= (0..7)
0 1 0 0 0 0 0 0
##$HPPRGN= 0
##$IN= (0..63)
0.0003244 0 0 0 0 0 0 0 0 0 0 0 0 0 0 0 0 0 0 0 0 0 0 0 0 0 0 0 0 0 0 0
```

```
0 0 0 0 0 0 0 0 0 0 0 0 0 0 0 0 0 0 0 0 0 0 0 0 0 0 0 0 0 0 0 0
##$INF= (0..7)
0 324.4 0 0 0 0 0 0
##$INP= (0..63)
0 0 0 0 0 0 0 0 0 0 0 0 0 0 0 0 0 0 0 0 0 0 0 0 0 0 0 0 0 0 0 0 0 0 0 0
0 0 0 0 0 0 0 0 0 0 0 0 0 0 0 0 0 0 0 0 0 0 0 0 0 0 0 0
##$INSTRUM= <spect>
##$INTEGFAC= (0..63)
0 0 0 0 0 0 0 0 0 0 0 0 0 0 0 0 0 0 0 0 0 0 0 0 0 0 0 0 0 0 0 0 0 0 0 0
0 0 0 0 0 0 0 0 0 0 0 0 0 0 0 0 0 0 0 0 0 0 0 0 0 0 0 0
##$L= (0..31)
1 1 1 1 1 1 1 1 1 1 1 1 1 1 1 1 1 1 1 1 1 1 1 1 1 1 1 1 1 1 1 1
##$LFILTER= 100
##$LGAIN= -5
##$LINPSTP= 0
##$LOCKED= yes
##$LOCKFLD= -1084
##$LOCKGN= 104.69896697998
##$LOCKPOW= 4
##$LOCKPPM= 4.69999980926514
##$LOCNUC= <2H>
##$LOCPHAS= 142.1351
##$LOCSHFT= yes
##$LOCSW= 0
##$LTIME= 0.349999994039536
##$MASR= 4200
##$MASRLST= <masrlst>
##$MULEXPNO= (0..15)
0 0 0 0 0 0 0 0 0 0 0 0 0 0 0 0
##$NBL= 1
##$NC= -6
##$NLOGCH= 5
##$NOVFLW= 0
##$NS= 16
##$NUC1= <1H>
##$NUC2= <13C>
##$NUC3= <15N>
##$NUC4= <off>
##$NUC5= <off>
##$NUC6= <off>
##$NUC7= <off>
##$NUC8= <off>
##$NUCLEUS= <off>
##$NUSLIST= <automatic>
##$NusAMOUNT= 50
##$NusFPNZ= no
##$NusJSP= 0
##$NusSEED= 54321
##$NusSPTYPE= 0
##$NusT2= 1
##$NusTD= 0
##$O1= 7793.69
##$O2= 23899.0843
##$O3= 11075.45013
##$O4= 0
##$O5= 0
##$O6= 4467.11
```

##$O7= 4467.11
##$O8= 4467.11
##$OVERFLW= 0
##$P= (0..63)
10.33 11.6 20.66 12 24 13.2 20 40 500 25 50 1000 2000 259 162 1375 800
2500 100000 600 2000 32 64 947 663 95 55 10.33 0 4000 160 512 0 0 0 101
0 0 0 1957.91 1431.72 1389 884 1074 126 72 48 1326 2084 63158 3789 407
884 0 0 821 500 1011 0 0 253 100 350 1500
##$PACOIL= (0..15)
0 0 0 0 0 0 0 0 0 0 0 0 0 0 0 0
##$PAPS= 0
##$PARMODE= 1
##$PCPD= (0..9)
0 55 50 220 0 0 0 0 0 0
##$PEXSEL= (0..9)
1 1 1 1 1 1 1 1 1 1
##$PHCOR= (0..31)
0 0 0 0 0 0 0 0 0 0 0 0 0 0 0 0 0 0 0 0 0 0 0 0 0 0 0 0 0 0 0 0
##$PHLIST= <>
##$PHP= 1
##$PH_ref= 0
##$PL= (0..63)
120 120 120 120 120 120 120 120 120 120 120 120 120 120 120 120 120 120
120 120 120 120 120 120 120 120 120 120 120 120 120 120 120 120 120 120
120 120 120 120 120 120 120 120 120 120 120 120 120 120 120 120 120 120
120 120 120 120 120 120 120 120 120 120
##$PLSTEP= 0.1
##$PLSTRT= -6
##$PLW= (0..63)
0 14 188 400 0 0 0 0 0 5.9757e-05 3.7348 0.49386 10.829 0 2.7072 43.315
8.4628 0 14 0.49386 5.2222 0.6484 11.75 64 0.048128 24.237 3.3437
45.385
28.456 0.42294 10.829 43.315 2.3903e-06 0 0 0 33.851 0 0 0 0 0 0 0 0 0
0 0 0 0 0 0 0 0 0 0 0 0 0 0 0 0 0 0
##$PLWMAX= (0..7)
74 321.3 1500 0 0 0 0 0
##$PQPHASE= 0
##$PQSCALE= 1
##$PR= 1
##$PRECHAN= (0..15)
-1 5 0 1 4 -1 -1 -1 -1 -1 -1 -1 -1 -1 -1 -1
##$PRGAIN= 0
##$PROBHD= <Z140930_0001 (CP TCI 950 H&F/C/N-D-05 Z)>
##$PULPROG= <b_trosy_hzdqc.cw>
##$PW= 0
##$PYNM= <>
##$ProjAngle= 0
##$QNP= 0
##$RD= 0
##$RECCHAN= (0..15)
0 2 1 0 0 0 0 0 0 0 0 0 0 0 0 0
##$RECPH= 0
##$RECPRE= (0..15)
-1 5 0 -1 -1 -1 -1 -1 -1 -1 -1 -1 -1 -1 -1 -1
##$RECPRFX= (0..15)
1 0 0 0 0 1 0 0 0 0 0 0 0 0 0 0
##$RECSEL= (0..15)

```
0 1 2 0 0 0 0 0 0 0 0 0 0 0 0 0
##$RG= 165.58
##$RO= 0
##$RSEL= (0..15)
0 1 2 0 3 0 0 0 0 0 0 0 0 0 0 0
##$S= (0..7)
83 83 83 83 83 83 83 83
##$SELREC= (0..9)
0 0 0 0 0 0 0 0 0 0
##$SFO1= 950.45779369
##$SFO2= 239.0147420843
##$SFO3= 96.31933745013
##$SFO4= 950.45
##$SFO5= 950.45
##$SFO6= 950.45446711
##$SFO7= 950.45446711
##$SFO8= 950.45446711
##$SOLVENT= <H2O+D2O>
##$SOLVOLD= <off>
##$SP= (0..63)
120 120 120 120 120 120 120 120 120 120 120 120 120 120 120 120 120 120
120 120 120 120 120 120 120 120 120 120 120 120 120 120 120 120 120 120
120 120 120 120 120 120 120 120 120 120 120 120 120 120 120 120 120 120
120 120 120 120 120 120 120 120 120 120
##$SPECTR= 0
##$SPINCNT= 0
##$SPNAM= (0..63)
<> <Sinc1.1000> <G4.256> <Q3.1000> <G4.256> <Q3.1000> <G4tr.256>
<Q3.1000>
<G4tr.256> <Q3_surbop.1> <Q5.1000> <Sinc1.1000> <Q5tr.1000>
<Crp80,0.5,20.1>
<Crp48,1.5,20.2> <Q3_surbop.1> <Q3_surbop.1> <Q3_surbop.1>
<Crp60_xfilt.2>
<Iburp2.1000> <Q3.1000> <Reburp.1000> <Pc9_4_90.1000> <Pc9_4_90.1000>
<Reburp.1000>
<Pc9_4_90.1000> <Reburp.1000> <Pc9_4_90.1000> <Eburp2.1000>
<Eburp2tr.1000>
<Bip720,50,20.1> <Crp48,1.5,20.2> <Gaus1_180i.1000> <Q3_surbop.1>
<Q3.1000>
<Reburp.1000> <0.0> <0.0> <Iburp2.1000> <Bip720,50,20.1> <Reburp.1000>
<Bip720,100,10.1> <> <> <> <> <> <> <> <> <> <> <> <> <> <> <> <> <> <>
<> <> <> <>
##$SPOAL= (0..63)
0.5 0.5 1 0.5 1 0.5 0 0.5 0 0.5 1 0.5 0 0.5 0.5 0.5 0.5 0.5 0.5 0.5 0.5
0.5 1 1 0.5 1 0.5 0 1 0 0.5 0.5 0.5 0.5 0.5 0.5 0.5 0.5 0.5 0.5 0.5 0.5
0.5 0.5 0.5 0.5 0.5 0.5 0.5 0.5 0.5 0.5 0.5 0.5 0.5 0.5 0.5 0.5 0.5 0.5
0.5 0.5 0.5 0.5
##$SPOFFS= (0..63)
0 0 0 0 0 0 0 0 0 0 0 0 0 0 0 0 0 0 0 0 0 0 0 0 0 0 0 0 0 0 0 0 0 0 0 0
0 0 0 0 0 0 0 0 0 0 0 0 0 0 0 0 0 0 0 0 0 0 0 0 0 0 0 0
##$SPPEX= (0..63)
0 0 0 0 0 0 0 0 0 0 0 0 0 0 0 0 0 0 0 0 0 0 0 0 0 0 0 0 0 0 0 0 0 0 0 0
0 0 0 0 0 0 0 0 0 0 0 0 0 0 0 0 0 0 0 0 0 0 0 0 0 0 0 0
##$SPW= (0..63)
0 0.0043077 140.63 179.82 140.63 179.82 140.63 179.82 140.63 16.97
10.159
0.00026923 10.159 55.151 26.472 28.456 16.97 16.97 18.924 6.0809 179.82
```

3.9144 0.0066594 0.03140509 0.5767664 0.049554 1.2005 0.049554 0.34773
0.34773 6.0224 6.6181 8.8436e-06 16.97 91.37 5.6635 0 0 0.87534 104.86
251.65 169.85 0 0 0 0 0 0 0 0 0 0 0 0 0 0 0 0 0 0 0 0 0 0
##$SUBNAM= (0..9)
<> <> <> <> <> <> <> <> <> <>
##$SW= 16.0384843390493
##$SWIBOX= (0..19)
0 1 2 4 0 0 0 0 0 0 0 0 0 0 0 0 0 0 0 0
##$SW_h= 15243.9024390244
##$SWfinal= 0
##$SigLockShift= 0
##$TD= 3072
##$TD0= 1
##$TD_INDIRECT= (0..7)
0 512 0 0 0 0 0 0
##$TDav= 1
##$TE= 276.9994
##$TE1= 278.0853
##$TE2= 0
##$TE3= 0
##$TE4= 0
##$TEG= 300
##$TE_MAGNET= 0
##$TE_PIDX= 1
##$TE_STAB= (0..9)
2 2 0 0 0 0 0 0 0 0
##$TL= (0..7)
120 120 120 120 120 120 120 120
##$TOTROT= (0..63)
0 0 0 0 0 0 0 0 0 0 0 0 0 0 0 0 0 0 0 0 0 0 0 0 0 0 0 0 0 0 0 0 0 0 0 0
0 0 0 0 0 0 0 0 0 0 0 0 0 0 0 0 0 0 0 0 0 0 0 0 0 0 0 0
##$TUBE_TYPE= <>
##$USERA1= <>
##$USERA2= <>
##$USERA3= <>
##$USERA4= <>
##$USERA5= <>
##$V9= 5
##$VALIDCODE= -1
##$VALIST= <>
##$VCLIST= <>
##$VDLIST= <>
##$VPLIST= <>
##$VTLIST= <>
##$WBST= 1024
##$WBSW= 20
##$XGAIN= (0..3)
0 0 0 0
##$XL= 0
##$YL= 0
##$YMAX_a= 62146
##$YMIN_a= -86300
##$ZGOPTNS= <>
##$ZL1= 120
##$ZL2= 120
##$ZL3= 120
##$ZL4= 120

```
##END=
```
